# Differential recognition of opioid analgesics by µ opioid receptors: Predicted interaction patterns correlate with ligand-specific voltage sensitivity

**DOI:** 10.1101/2021.12.02.470941

**Authors:** Sina B. Kirchhofer, Victor Jun Yu Lim, Julia G. Ruland, Peter Kolb, Moritz Bünemann

## Abstract

The µ opioid receptor (MOR) is the key target for analgesia, but the application of opioids is accompanied by several issues. There is a wide range of opioid analgesics, differing in their chemical structure and their properties in receptor activation and subsequent effects. A better understanding of ligand-receptor interactions and resulting effects is important. Here, we calculated the respective binding modes for several opioids and analyzed fingerprints of ligand-receptor interactions. We further corroborated the binding modes experimentally by cellular assays. As ligand-induced modulation of activity due to changes in membrane potential was displayed by MOR, we further analyzed the effects of voltage sensitivity of this receptor. With a combined in silico and in vitro approach, we defined discriminating interaction patterns for the ligand-specific voltage sensitivity. With this, we present new insights for interactions likely in ligand recognition and their specific effects on activation of the MOR.

## Introduction

Opioids, agonists at the µ opioid receptor (MOR), are the most effective analgesics in clinical use. However, their pain killing effects are accompanied by severe side effects, like respiratory depression and addiction. Their high risk for abuse and overdose led to the opioid crisis in the USA with nearly 50.000 deaths caused by opioid-overdose in 2019 alone, on a rising trend (CDC, 2021). Especially synthetic drugs, such as fentanyl, are responsible for the majority of the observed deaths. The currently used opioid analgesics differ not only in their chemical structure, but also with respect to their potency, efficacy and kinetics to activate Gi/o proteins via MOR. Furthermore, they may exhibit differences in their efficacy to induce arrestin recruitment to MOR. There are already attempts to develop more effective and safer opioids through a structure-based approach, shown by the identification of the so-called biased opioids (Manglik et al., 2016; Schmid et al., 2017) which show a bias towards G protein activation versus arrestin recruitment. Whether or not a G protein bias would be beneficial with respect to minimized side effects is disputed (Gillis et al., 2020; Kliewer et al., 2020; Raehal et al., 2005). However, based on the above-mentioned differences between the different opioids, it is important to understand details of ligand-receptor interactions. We recently showed that ligand-induced MOR activity was modulated by the membrane potential, and the effect and extent of this voltage sensitivity was ligand specific (Ruland et al., 2020). Since the first report of voltage sensitivity of the muscarinic M_2_ receptor (Ben-Chaim et al., 2003), several other GPCRs have been observed to be modulated in their activity depending on the membrane potential. Moreover, these effects were found to be ligand specific (Birk et al., 2015; López-Serrano et al., 2020; Moreno-Galindo et al., 2016; Navarro-Polanco et al., 2011; Rinne et al., 2015, 2013), indicating the voltage effect on GPCRs is influenced by the receptor-ligand interaction. However, a general mechanism of voltage sensitivity is still elusive. The expression of the MOR in highly excitable tissue, such as neurons, makes this receptor an interesting candidate for further analysis of the voltage sensitivity. Moreover, due to the clinical relevance of opioids, a wide range of ligands of the MOR have been described. Analysis of the interactions of these ligands with the receptor in general would not only give new information on molecular determinants of ligand-specific voltage sensitivity, these new insights could also be used in the fine tuning of new safer and more effective opioids. Therefore, we analyzed the predicted binding modes of many opioids, detected important key interactions and interaction groups and correlated these with the effects voltage has on the MOR. To do so, we performed molecular docking calculations for 10 opioid ligands, including the clinically most relevant ones. Subsequently, we experimentally corroborated the predicted binding modes by Förster resonance energy transfer (FRET) based assays in HEK293T cells. The analysis of the ligand-specific voltage sensitivity of the MOR was further performed with FRET-based functional assays under direct control of the membrane potential, revealing a correlation of the particular binding mode of the ligand and the specific voltage sensitivity of the MOR. Based on these observations, by means of site-directed mutagenesis, we identified receptor regions determining the voltage effect on the MOR.

## Results

### Voltage sensitivity of the MOR is ligand specific

Voltage sensitivity of the MOR was investigated by utilizing single cell FRET assays to study G protein activity as well as recruitment of arrestin3 to the MOR under conditions of whole cell voltage clamp. To detect the effect of voltage on G protein activity, HEK293T cells were transfected with wild-type µ opioid receptors and Gα_i_-mTurquoise, cpVenus-Gγ_2_ and Gβ, in order to monitor G_i_ protein activity by a decrease in the FRET emission ratio (van Unen et al., 2016). Agonists were applied at concentrations close to the EC_50_-value to avoid signal saturation. The level of maximal stimulation was determined by application of a saturating concentration of DAMGO in all FRET recordings. Application of morphine at -90 mV induced a robust Gα_i_ activation (Fig. 1A), depolarization to +30 mV enhanced the Gα_i_ activation strongly and the effect was reversible after repolarization. A similar protocol was applied to cells stimulated with methadone (Fig. 1B) or fentanyl (Fig. 1C). Here, however, the depolarization-induced a decrease in Gα_i_ activation. Voltage affected only the FRET signal if a ligand was present and MOR was expressed (Fig. S1A, B-G). Ligand dependence of the voltage sensitivity, mainly based on a change in efficacy in receptor activation, was additionally reported previously for morphine, Met-enkephalin, DAMGO and fentanyl (Ruland et al., 2020). Therefore, the MOR shows a strong ligand-specific voltage sensitivity.

**Figure 1:**
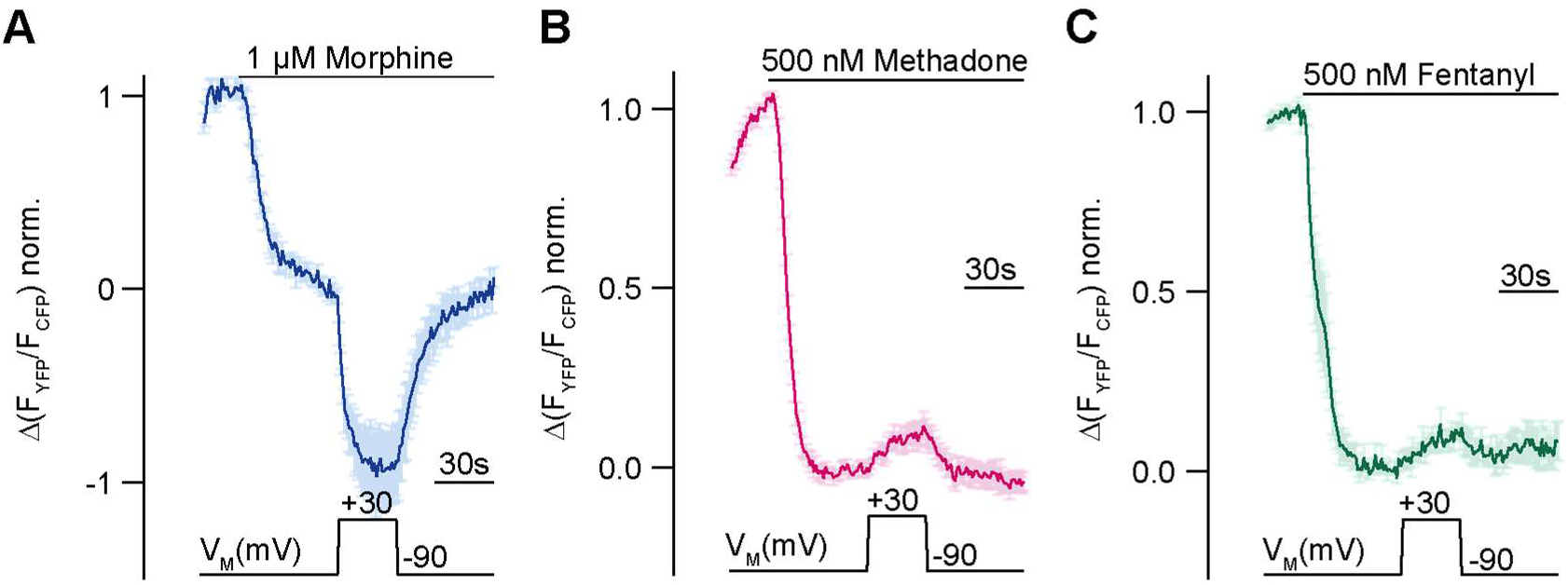
Voltage sensitivity of the MOR is ligand specific. **(A-C)** Averaged FRET-based single cell recordings of MOR-induced Gα_i_ activation under voltage clamp conditions with WT receptor, Gα_i_- mTurquoise, cpVenus-Gγ_2_ and Gβ in HEK293T cells are plotted for the indicated agonists (mean ± SEM; A: n=8, B: n=13, C: n=12). The applied voltage protocol is indicated below.

### Binding modes of different opioids at the MOR reveal differential interaction patterns

To gain mechanistic insights into this ligand-specific voltage sensitivity, we evaluated the binding modes of several opioid ligands by molecular docking. Our docking calculations were performed based on the crystal structure of the active MOR (PDB: 5C1M (Huang et al., 2015)). We decided not to use the cryo-EM structure of the MOR bound to the G proteins (PDB: 6DDE (Koehl et al., 2018)), as the resolution was lower and an alignment of both structures (Fig. S2A) displayed only minor differences in the ligand binding site (mass-weighted RMSD: 1.21A of residues in pocket). The docking calculations revealed different binding modes for the different opioids. The binding mode for morphine (Fig. 2A) suggested D147^3.32^, Y148^3.33^ and both water molecules between helices 3 and 5 as important interaction partners, and M151^3.36^, V236^5.42^, H297^6.52^, W293^6.48^, V300^6.55^ and Y326^7.43^ as possible interactions, as well (numbers in superscript are according to the Ballesteros-Weinstein enumeration scheme for GPCRs (Ballesteros and Weinstein, 1995)). In contrast, the binding mode for methadone (Fig. 2B) indicated only a salt bridge with D147^3.32^, and hydrophobic interactions and/or aromatic-aromatic stacking interactions with V236^5.42^, H297^6.52^, W293^6.48^ and Y326^7.43^. In contrast, fentanyl (Fig. 2C) was predicted to form only one H-bond with Y326^7.43^ via its amide carbonyl. Indeed, the charged amine was not predicted to form an ionic interaction with the conserved D147^3.32^. In addition, N127^2.63^, D147^3.32^, C217^45.50^ and H297^6.52^ were possible interactions for fentanyl. As the D147^3.32^ interaction was known to be important for opioids (Surratt et al., 1994), and the formation of a salt bridge between the D147^3.32^ and fentanyl was already postulated (Weltrowska et al., 2010), we took a closer look at the pose of fentanyl. As the docking calculation always preferred a conformation with the nitrogen-bound hydrogen in our hands, we explored a different path. First, we inverted the orientation of the hydrogen and the propyl-benzene group at the formally charged nitrogen. Then, we exhaustively sampled the three rotatable bonds of the propyl- benzene group of fentanyl with Pymol at five-degree intervals, resulting in 186.624 poses. Each pose was minimized with OEtoolkit with the steepest-descent algorithm and poses were ranked based on their final force-field energy. The best scored pose (Fig. 2D) indeed displayed a salt bridge with D147^3.32^. Encouragingly, top 10.000 poses were quite similar in orientation, with only two clusters discernable (Fig. S2B). Clustering of the top 10.000 poses and the spread of energy values after minimization is shown in figure S2C.The binding modes of other opioids can be found in figure S2D-J. All binding modes were further investigated with a fingerprint analysis, a computational evaluation converting the interactions between a ligand and the receptor into a string of numbers, i.e., a vector. The interaction patterns (Fig. 2E) that contribute strongest to the first two principal components emerged from this analysis and are plotted in Fig. 2F. On one side, interactions defining the first principal component (PC1, describing 28% of the variance observed in the interactions) could be found within helices 5 and 6 (V236^5.42^, H297^6.52^ and V300^6.55^). On the other side, key interactions defining the second principal component (PC2, describing 14 % of the variance observed in the interactions) were mostly found in extracellular loop 1 (W133^23.50^), helix 3 (D147^3.32^ and Y148^3.33^) and helix 6 (I296^6.51^). Residues deeper in the binding pocket were found to be important for both components (M151^3.36^, W293^6.48^ and Y326^7.43^). The principal component analysis therefore revealed diverse interaction patterns of the different opioid ligands with MOR. The calculated fingerprint of the docked fentanyl and the best-scored fentanyl conformer were nearly identical (Fig. 2F). The fingerprints of the second and third best scored conformers (Fig. S2L-M) were comparable as well. We further excluded our reference agonist DAMGO from this analysis, as we were not able to calculate a reliable binding mode for DAMGO due to the high flexibility of this peptidergic ligand. Analysis of the fingerprint of the published binding mode of DAMGO (Koehl et al., 2018) revealed a completely different interaction pattern (Fig. S2K) in comparison to the other opioids, putting it outside of the applicability domain. This is likely due to larger size of the peptide DAMGO in comparison to the non-peptide opioid agonists.

**Figure 2:**
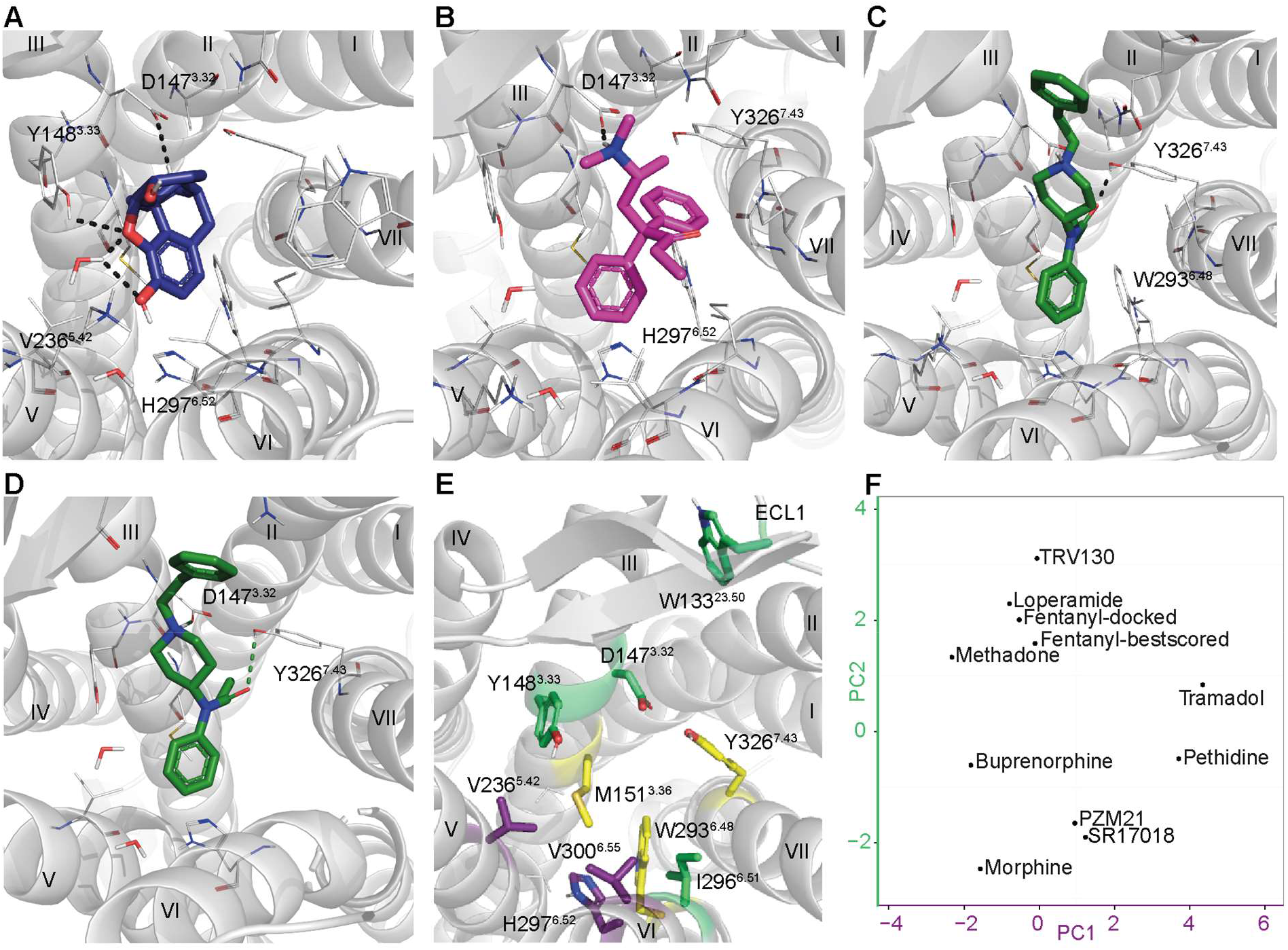
Predicted binding modes of different opioids at the MOR reveal differential interaction patterns. **(A-C)** Binding modes of morphine (A), methadone (B) and fentanyl (C) docked to the MOR are illustrated as a view from the extracellular side, H-bonds are indicated as dotted line. **(D)** Binding mode of the best scored fentanyl conformer. **(E)** Important interactions of several opioid ligands docked to MOR were identified by a fingerprint analysis which led to the definition of the principal components plotted in (F). Interactions contributing strongest to component 1 (PC1) can be found within helices five and six (V236^5.42^, H297^6.52^ and V300^6.55^, depicted in violet), whereas important interactions contributing strongest to component 2 (PC2) are mostly found in extracellular loop 1, and helices 3 and 6 (W133^23.50^, D147^3.32^, Y148^3.33^ and I296^6.51^, depicted in green). Residues depicted in yellow (M151^3.36^, W293^6.48^ and Y326^7.43^) are important interactions for both components. **(F)** Analyzed agonists were plotted with respect to the contribution of PC1 versus PC2 for their binding based on the fingerprint analysis.

### Functional effects of site-directed mutagenesis support calculated binding modes of different opioids at the MOR

To experimentally corroborate the observed docking poses, we performed site-directed mutagenesis of several residues that were predicted to be important or not in the binding pocket of the MOR. These mutations were first introduced in silico, and we repeated the docking for each ligand with the mutated receptor, analyzed the new binding mode to make a prediction about the likely effect of the mutation on binding and then evaluated the effects in functional assays in living cells. We identified Y148^3.33^ as one of the main interaction partners for morphine (Fig. 2A). The replacement of this residue by F (and with this, exclusion of the two water molecules from the docking calculation, as the F would prohibit this interaction) led to a changed binding mode for morphine (Fig. 3A), with now only the H-bond to D147^3.32^ left as interaction. We did not mutate D147^3.32^, as it had already been determined as the most important amino acid for ligand binding and activation of MOR previously (Surratt et al., 1994). Next, we determined concentration-response curves for G protein activation in single-cell FRET measurements for the different modified receptors. To that end, we measured Gα_i_ activation induced by MOR WT (Fig. 3B) or the mutated version of MOR (Fig. 3C) at different concentrations of morphine and compared it to the maximal activation by DAMGO. We plotted these as concentration-response curves (Fig. 3D) and calculated EC_50_-values for each receptor variant. The mutation of Y148^3.33^, V236^5.42^ and H297^6.52^, respectively, led to a strong right shift of the EC50-value for morphine-induced Gαi activation (more than 4 orders of magnitude), indicating the importance of these residues for proper morphine binding. For methadone, we identified H297^6.52^ as important interaction (Fig. 2B), replacement by an A resulted in the formation of an H-bond with Y148^3.33^ (Fig. 3E). This change in the predicted binding mode was consistent with by the right shift of EC_50_-value for Gα_i_ activation by 4 orders of magnitude (Fig. 3F). Furthermore, the identification of Y326^7.43^ as important interaction was verified by a right shift in the EC_50_-value for Gα_i_ activation of 2 orders of magnitude (Fig. 3F). The same could be seen for fentanyl (Fig. 3 G-H), as the EC_50_-value displayed a rightward shift by 5 orders of magnitude. Residue W293^6.48^, part of the CWxP motif, which is known to be important for the activation of class A GPCRs (Shi et al., 2002), was identified as important interaction for both methadone and fentanyl as well. Replacement by the smaller F resulted in an altered pose for fentanyl (Fig. S3W). This was strongly supported by the nearly completely abolished Gα_i_ activation for this mutation (Fig. 3H). In contrast, for methadone we didn’t observe a changed binding pose (Fig. S3X), and the Gα_i_ activation was increased in good agreement (Fig. S3J, left shift by 2 orders of magnitude). To give an overview of all mutations and their influence on Gα_i_ activation (Fig. S3A-R), we plotted all measured EC_50_-values normalized to the WT receptor in a heatmap (Fig. 3I). This experimentally corroborated the binding modes we observed in the docking calculations based on changes in potency of receptor activation. We did not evaluate the influence on efficacy of receptor activation of the receptor mutants (Fig. S3S-V).

**Figure 3:**
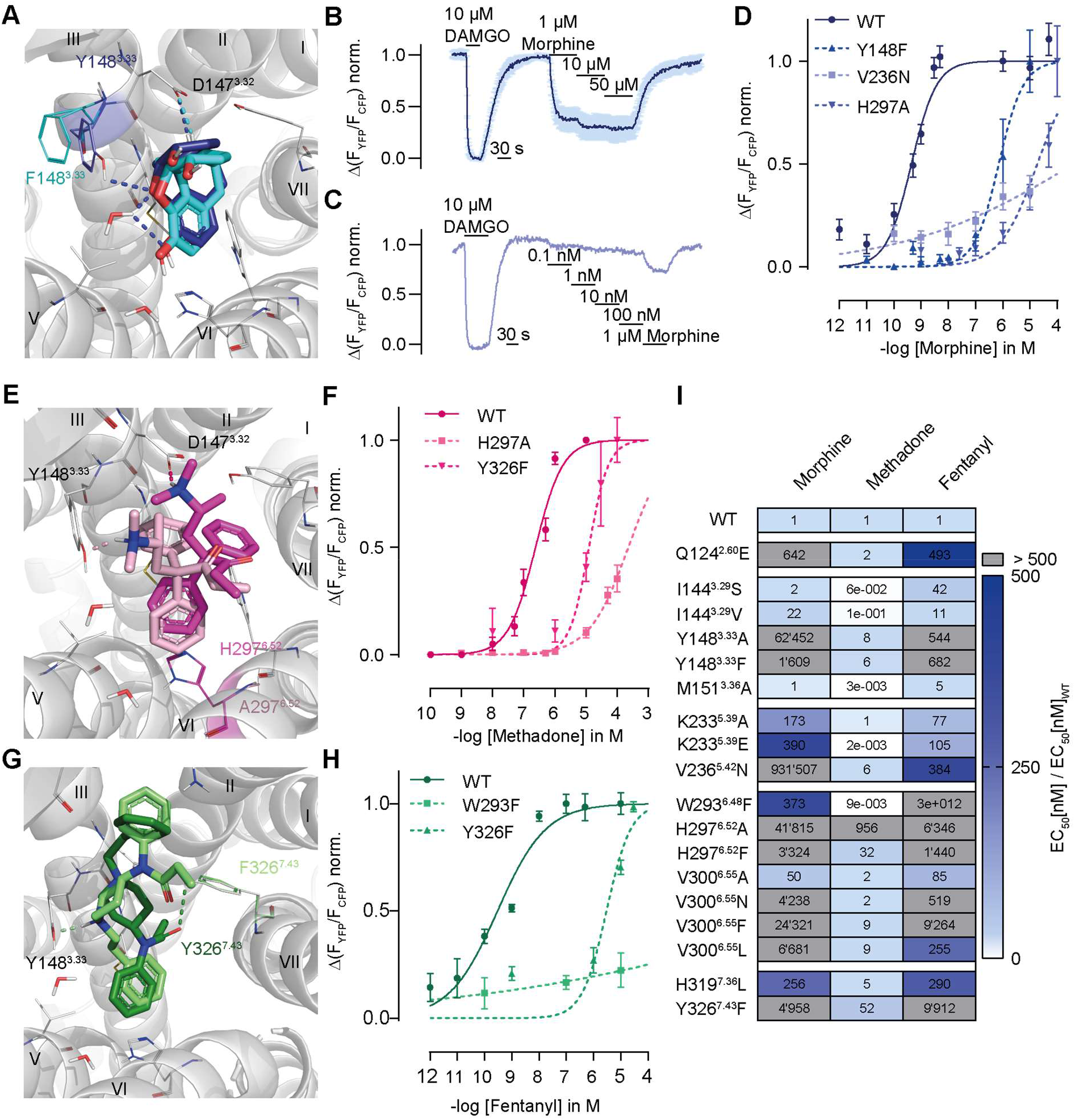
Functional effects of point mutations corroborate binding poses of different opioids at the MOR. **(A, E, G)** Binding modes of morphine (A; WT depicted in dark blue, Y148^3.33^F in cyan), methadone (E, WT in dark pink, H297^6.52^A in magenta) and fentanyl (G, WT in dark green, Y326^7.43^F in bright green) docked to MOR are illustrated as in Fig. 2. The insertion of a point mutation changed these binding modes. **(B, C)** FRET-based single cell recordings of MOR-induced Gα_i_ activation for MOR WT (B, mean ± SEM, n=6) and MOR-Y148^3.33^F (C) were used for the fit of concentration-response curves. Maximum activation for normalization was induced by 10 µM DAMGO. **(D, F, H)** Representative concentration-response curve for Gα_i_ activation induced by the depicted agonist were fitted for WT and mutated versions of MOR. For simplification, maximum Gα_i_ activation was set to 1, the EC_50_-values (D: morphine, WT=0.45 nM, Y148^3.33^F=716 nM, H297^6.52^A=19 µM, V236^5.42^N=415 µM; F: methadone, WT=238 nM, Y326^7.43^F=12 µM, H297^6.52^A=228 µM; H: fentanyl: WT=0.3 nM, Y326^7.43^F= 3 µM, W293^6.48^F= >1mM) were calculated, normalized to WT and plotted in (I). (I) The normalized EC_50_-values for Gα_i_ activation were plotted in a heatmap showing the loss in EC_50_-value depending on the point mutation.

### Binding mode is consistent with agonist specific voltage sensitivity of the MOR

As we saw different patterns in the predicted binding modes of the opioid ligands, we examined these ligands for their voltage sensitivity by analyzing the extent and direction of the effect of depolarization on Gα_i_ activation (Fig. S4). We compared the effects between the ligands (Fig. 4A), with the response at +30 mV normalized to the response at -90 mV. For this, we applied the agonist at a suitable concentration to induce a robust and equivalent Gα_i_ activation level in comparison to DAMGO (Fig. S4A). This led to a great variance of the direction and magnitude of voltage-induced effects, depending on the opioid ligand used for stimulation of Gα_i_ activation. Buprenorphine (Fig. S4A) and pethidine (Fig. S4B) enhanced their Gα_i_ activation strongly due to depolarization, comparable to morphine (Fig. 3A). In contrast, DAMGO (Fig. S4C), tramadol (Fig. S4D) and PZM21 (Fig. S4E) induced a slightly enhanced Gα_i_ activation. SR17018 (Fig. S4F) showed no apparent voltage sensitive behavior. Moreover, loperamide (Fig. S4G) and TRV130 (Fig. S4H) showed a voltage dependent decrease in Gα_i_ activation, comparable to the effect of fentanyl. Thus, opioid ligands can be grouped according their direction of voltage sensitivity. Comparing the docking poses of the opioids and their analyzed fingerprints, it becomes obvious, that the voltage sensitivity of agonists is correlated to the predicted ligand-receptor interaction pattern, as defined by the fingerprint analysis (Fig. 4B). Further analysis of the main interactions of the two groups separately resulted in the identification of distinct interaction motifs for both groups (Fig. 4C). The ligands that showed enhanced activity upon depolarization mainly interacted with ECL2 (T218^45.51^ and L219^45.52^) and helix 6 (W293^6.48^, H297^6.52^ and V300^6.55^), while the ligands exhibiting a decrease in activation upon depolarization mainly interact with helix 3 (I144^3.29^, Y148^3.33^ and M151^3.36^) and ECL2 (C217^45.50^) in these calculations. Overlaying this information on the binding pocket, three separate main interaction regions or motifs can be discerned (Fig. 4C), marked in blue (important for depolarization-induced activation), red (important for depolarization-induced deactivation) and yellow (important for all ligands), correlating with the voltage sensitive behavior of the ligand. We excluded SR17018 and DAMGO from this analysis: while SR17018 showed a completely different binding mode (Fig. S2K), DAMGO’s pose resulted in a completely different fingerprint (Fig. S2B).

**Figure 4:**
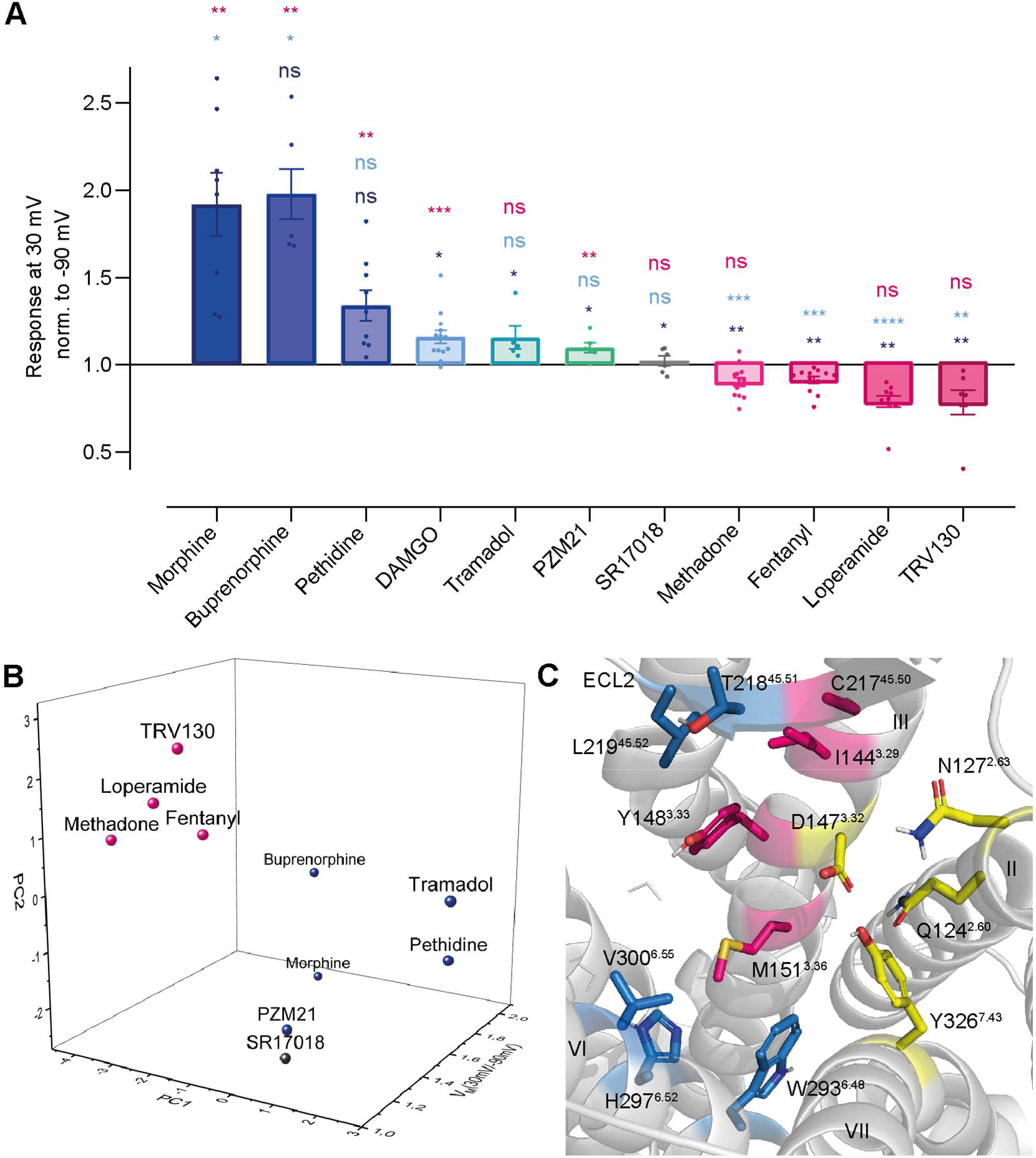
Predicted binding mode correlates with agonist specific voltage sensitivity of MOR. **(A)** FRET-based single cell recordings of Gα_i_ activation under voltage clamp conditions induced by different opioid agonist were analyzed for agonist specific voltage sensitive behavior. For this, the response of agonist-induced Gα_i_ activation at +30 mV was normalized to the response at -90 mV. The applied agonist concentrations induced approximately the same Gα_i_ activation level for all used agonists. Statistical significance was calculated compared to depolarization effect induced by morphine (dark blue), DAMGO (bright blue) and fentanyl (magenta) by an ordinary one-way ANOVA (p<0.0001) with Dunnett’s T3 multiple comparisons test (ns p>0.05, * p<0.05, ** p<0.005, *** p<0.0005). **(B)** Fingerprint analysis (Fig. 2E) was combined with the effects voltage displayed on the agonists and plotted as 3D plot. The agonists fell into groups with a group arrangement comparable to voltage sensitive effect, with morphine, buprenorphine, pethidine, tramadol and PZM21 in the group activating upon depolarization (blue) and methadone, fentanyl, loperamide and TRV130 deactivating upon depolarization (magenta). SR17018 showed no voltage sensitivity and also showed a different binding mode compared to the other agonists (black). **(C)** Detailed analysis of fingerprints split into groups regarding their voltage sensitive behavior resulted in the possibility to define the main predicted interaction partners for both groups. The group showing increased activation induced by depolarization mainly interacts with ECL2 (T218^45.51^ and L219^45.52^) and helix 6 (W293^6.48^, H297^6.52^ and V300^6.55^), depicted in blue. The group showing decreased activation induced by depolarization mainly interacts with ECL2 (C217^45.50^) and helix 3 (I144^3.29^, Y148^3.33^ and M151^3.36^), depicted in magenta. Residues depicted in yellow are interacting with both of the groups (N127^2.63^, Q124^2.60^, D147^3.32^ and Y326^7.43^).

### Voltage effect confirms the binding mode of ligands

Analysis of the voltage sensitivity of Gα_i_ activation induced by meptazinol (Fig. 5A) revealed a strong decrease in Gα_i_ activation upon depolarization (Fig. S5D). However, our docking analysis resulted in two slightly different possible binding modes (Fig. S5A-B) with two possible fingerprints (Fig. 5B), differently grouping with the two groups that we defined based on the different direction of the voltage effect. To test whether the binding mode depicted in Fig. S5A indeed was congruent with the voltage sensitivity of meptazinol, we measured concentration-response curves for Gα_i_ activation with mutations of the interacting residues (Fig. 5C). We corroborated the corresponding binding mode (Fig. S5A) by the exchange of N127^2.63^, the crucial interaction for this binding mode, resulting in a right shift of the concentration-response curve (Fig. 5C) and an EC_50_-value approx. 20 times higher than WT. As C217^45.50^, the crucial interaction for the alternative binding mode (Fig. S5B), is necessary for the formation of disulfide bridges within the receptor and a mutation would therefore destroy the receptor, we weren’t able to exchange this residue.

**Figure 5:**
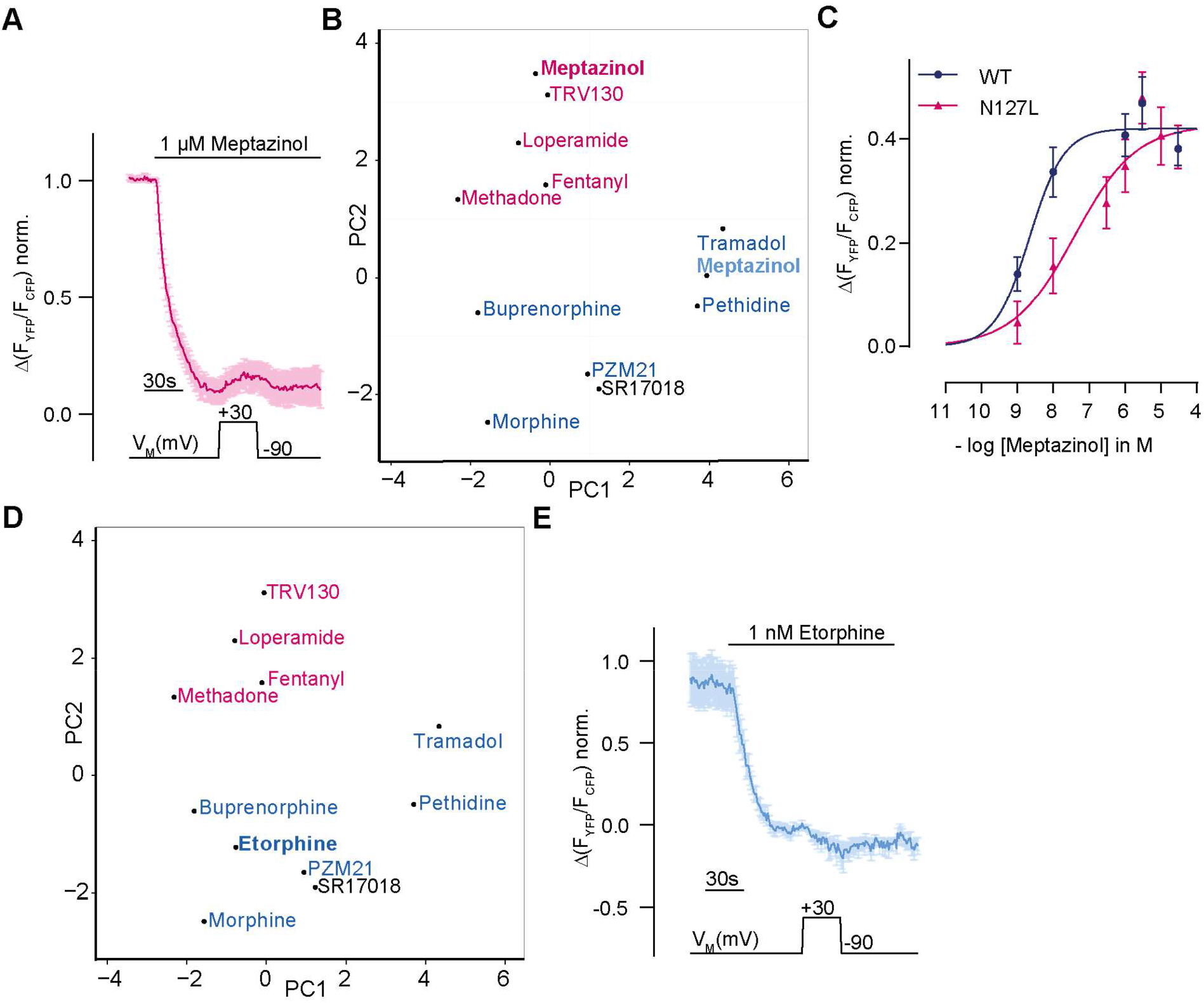
Voltage effect confirms the binding mode of meptazinol and binding mode of etorphine predicts voltage effect. **(A, E)** Average FRET-based single cell recordings of MOR-induced Gα_i_ activation under voltage clamp conditions are plotted for the indicated agonist (A: meptazinol, mean ± SEM, n=7, E: etorphine, mean ± SEM, n=8). The applied voltage protocol is indicated below. **(B)** Meptazinol was analyzed regarding its fingerprints (as shown in figure 2) resulting in two possible binding modes (Fig. S5 A and B) and two possible fingerprints which either group with the ligands showing activation upon depolarization (blue) or with the ligands showing deactivation upon depolarization (magenta). **(C)** Concentration-response curves for Gα_i_ activation induced by meptazinol, measured by single cell FRET, with MOR WT (dark blue) and mutated version of MOR were fitted and EC_50_-values were calculated (WT=2 nM and N127^2.63^L=39 nM). **(D)** Etorphine was docked (Fig. S5C) and analyzed regarding its fingerprint, resulting in a fingerprint grouping with the ligands showing activation upon depolarization.

### Docking-derived binding mode predicts voltage effect of ligands

We were not only able to confirm binding modes of ligands by the voltage effect, we were also able to do this vice versa by the prediction of the voltage effect of a certain ligand by analysis of its predicted binding mode. The binding mode of etorphine (Fig. S5C) was comparable to the binding modes of morphine (Fig. 2A) and buprenorphine (Fig. S2D), and the fingerprint grouped well with these agonists, which are displaying an activation upon depolarization (Fig. 5D). Based on this observation, we predicted that etorphine would display a voltage sensitivity in the same direction. This was confirmed by the analysis of voltage sensitivity of etorphine, revealing an increased Gα_i_ activation upon depolarization (Fig. 5E and S5D).

### Altered binding modes influence agonist-specific voltage sensitivity of the MOR

As we already showed that site-directed mutagenesis alters the predicted binding mode of the ligands, we evaluated the influence of these altered binding modes on the agonist-specific voltage sensitivity of the MOR to gain more information on potential molecular determinants for the voltage sensitivity. Therefore, we measured mutated MOR-induced Gα_i_ activation under voltage clamp conditions and compared this to the WT behavior. Agonists were applied in a concentration inducing comparable Gα_i_ activation levels, which were determined respectively (Fig. 3I and S3A-R). Due to this extensive evaluation of each mutation, it was not necessary to evaluate expression levels for each mutant. The mutation of Y148^3.33^ to F resulted in a reduced voltage effect of morphine (Fig. 6A), but still to an increased Gα_i_ activation upon depolarization. The insertion of an A at this position instead led to a strongly increased Gα_i_ activation, even stronger than the one for WT (Fig. 6A). The exchange of H297^6.52^ to an A increased the effect of depolarization on Gα_i_ activation induced by fentanyl with an even stronger decrease in Gα_i_ activation (Fig. 6B). It further changed the direction of voltage effect for methadone (Fig. 6C), now showing an increased Gα_i_ activation upon depolarization. The exchange of H297^6.52^ to an F led to a voltage effect of methadone comparable to WT behavior (Fig. 6C). The insertion of an F instead of Y326^7.43^ changed the direction of voltage effect for methadone (Fig. 6D). All effects on voltage sensitive behavior induced by the altered binding modes were plotted in a heatmap (Fig. 6E, based on data of Fig. S6A-C), where the agonist-induced response at +30 mV was normalized to the response at -90 mV. We did not analyze the effect of double-mutants, as these displayed only weak and not evaluable Gα_i_ activation (Fig. S6D). Overall, although the suggested binding mode of morphine changed, depolarization always increased Gα_i_ activation to a varying extent. For methadone and fentanyl, the altered predicted binding mode was consistent with the change in direction of the voltage effect of methadone- or fentanyl-induced Gα_i_ activation, which was now increasing upon depolarization in some cases. Overall, the strongest effects were induced by mutation of Y148^3.33^ and H297^6.52^. As already shown by the fingerprint analysis (Fig. 4C), the interactions with helices 3 and 6 seem to have the largest influence on voltage sensitivity, probably because of the largest voltage-induced movements upon depolarization. Especially mutations leading to more space close to these helices (Y148^3.33^A, M151^3.36^A and H297^6.52^A) shifted the voltage effect of the different agonists into enhanced depolarization-induced activation.

**Figure 6:**
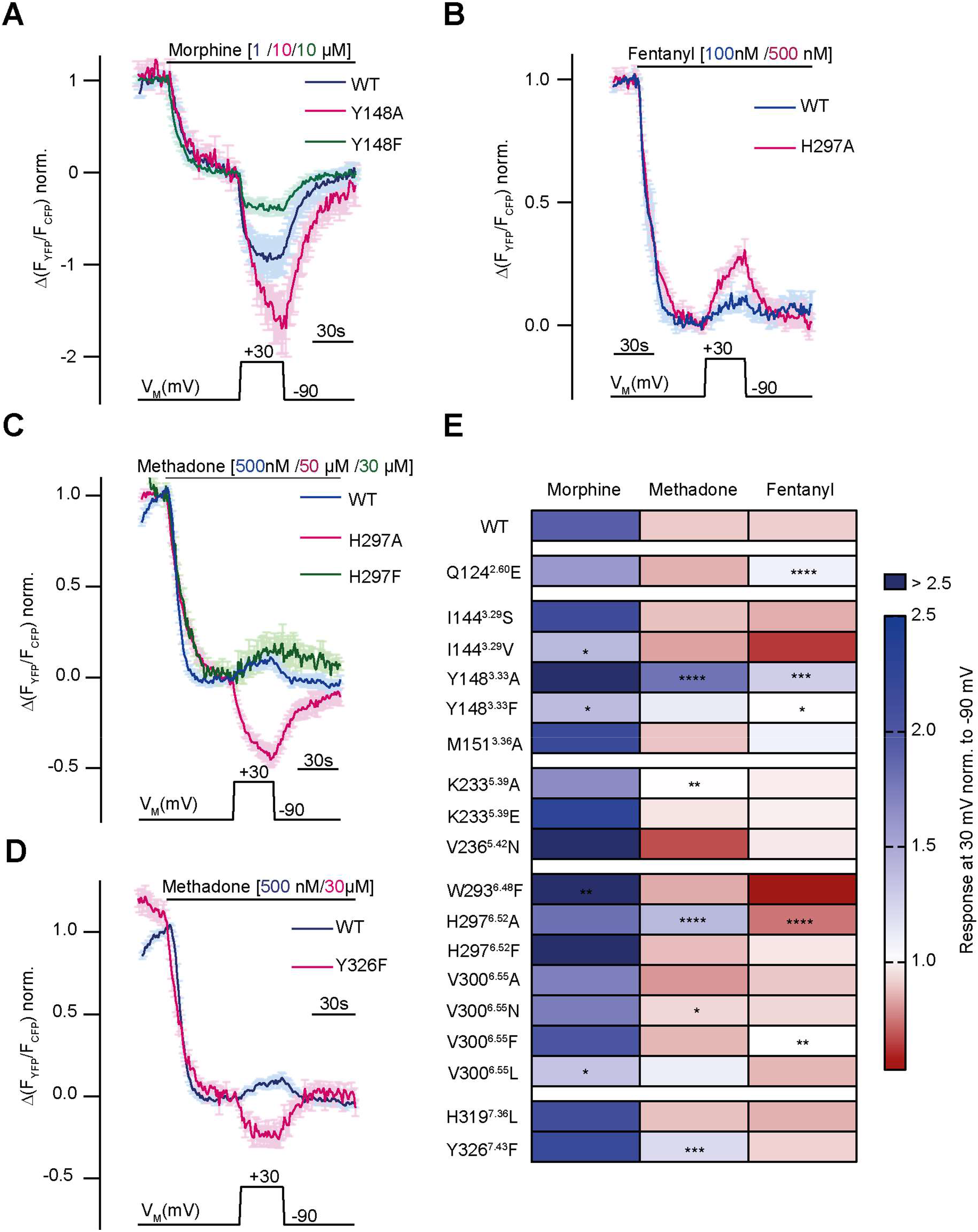
Altered binding modes influence agonist specific voltage sensitivity of the MOR. **(A-D)** Average (mean ± SEM) FRET-based single cell recordings of Gα_i_ activation measured in HEK293T cells under voltage clamp conditions are plotted for the indicated agonist and mutation (A: MOR WT (blue, n=8), MOR-Y148^3.33^A (magenta, n=9), MOR-Y148^3.33^F (green, n=6); B: MOR WT (blue, n=12), MOR- H297^6.52^A (magenta, n=10); C: MOR WT (blue, n=13), MOR-H297^6.52^A (magenta, n=11), H297^6.52^F (green, n=6); D: MOR WT (blue, n=13), MOR-Y326^7.43^F (magenta, n=6)). The applied voltage protocol is indicated below. **(E)** The analyzed depolarization effects on Gα_i_ activation induced by mutations were plotted in a heatmap regarding the applied agonist (applied concentrations induced for all used agonist the approx. same Gα_i_ activation level). Response of agonist-induced Gα_i_ activation at +30 mV was normalized to response at -90 mV, a value smaller 1 indicates a decreased Gα_i_ activation induced by depolarization (depicted in red), a value larger than 1 indicates an increased Gα_i_ activation induced by depolarization (depicted in blue), no voltage effect is indicated by a value around 1 (depicted in white). Significance was calculated compared to depolarization effect of the WT receptor and the respective agonist (unpaired t-test with Welch’s correction (ns p>0.05, * p<0.05, ** p<0.005, *** p<0.0005, **** p<0.0001)).

### Depolarization converts the antagonist naloxone to an agonist

Naloxone is the classical antagonist for the MOR. We analyzed the binding mode of naloxone by molecular docking, and, as the chemical structure of naloxone contains the morphinan scaffold and is highly related to morphine overall, we compared their predicted binding modes (Fig. 7A). Their binding modes were highly comparable, as expected due to their conserved scaffold. Only the direct interaction with Y148^3.33^ seems to be missing for naloxone. According to the fingerprint analysis naloxone belongs to the group of ligands that shows increased activation upon depolarization (Fig. 7B). As it was reported before that depolarization can convert GPCR antagonists to agonists (Gurung et al., 2008), we also evaluated naloxone with respect to voltage sensitivity. For this, we measured MOR-induced Gα_i_ activation under voltage clamp conditions. Application of naloxone at -90 mV induced no Gα_i_ activation (Fig. 7C), depolarization to +30 mV led to Gα_i_ activation up to a level of approx. 30% of the Gα_i_ activation induced by a saturating concentration of DAMGO. This effect was reversible after repolarization. We further analyzed this voltage effect by the application of different membrane potentials (Fig. S7A) and fitted these to a Boltzmann function (Fig. 7D). A comparison with the effects evoked by morphine in the same setting revealed that the net charge movements upon change in membrane potential, represented as z-values, were comparable with 1.17 for naloxone and 0.8 for morphine. Both values are also in the same range of z-values previously published for other GPCRs (Ben-Chaim et al., 2006; Birk et al., 2015; Kurz et al., 2020; Navarro-Polanco et al., 2011; Rinne et al., 2013). The half-maximal effective membrane potential for naloxone (V_50_: +31 mV) was shifted to a more positive V_M_ in comparison to morphine (V_50_: -29 mV), indicating that the conversion of naloxone from an antagonist to an agonist requires more positive membrane potential. We performed the identical analysis also for Gα_o_ activation (Fig. S7B), resulting in nearly identical V_50_ and z-values for data fitted to a Boltzmann function (Fig. S7C). Further, we checked if this effect is also visible in assays that show no amplification. For this, we measured the direct interaction of MOR-sYFP and arrestin3-mTur2 (Fig. S7D, see also (Ruland et al., 2020)) under voltage clamp conditions. In this case, naloxone induced no arrestin recruitment to the receptor, neither at -90 mV nor at +45 mV. This was comparable to effects of weak partial agonists like tramadol, which induce no arrestin recruitment as well (Fig. S7E). In order to further verify the observed voltage-induced conversion from antagonist to agonist for naloxone, we measured MOR-evoked inward GIRK currents at different holding potentials, as previously described (Ruland et al., 2020). We applied naloxone and compared the evoked K^+^ current to a saturating concentration of DAMGO (Fig. 7E) at -90 mV and -20 mV. The response evoked by naloxone at -90 mV was approx. 8% of the response evoked by DAMGO, whereas the response at -20 mV was approx. 16% of the response evoked by DAMGO (Fig. 7F), indicating a significantly increased naloxone-induced current at -20 mV. To verify that the measured currents were K^+^ currents, we applied Ba^2+^ before and after every agonist or antagonist application.

**Figure 7:**
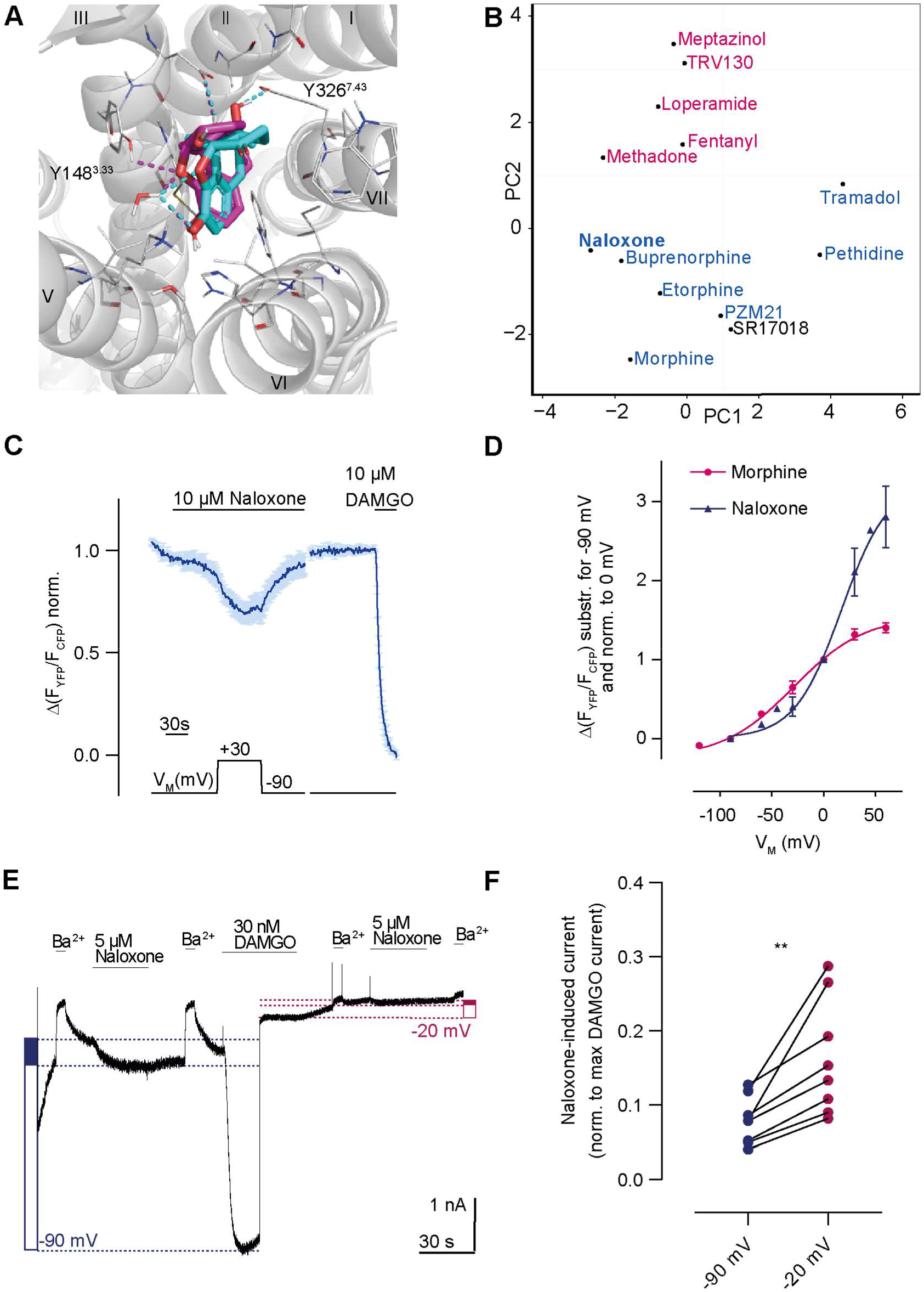
Depolarization converts the antagonist naloxone to an agonist. **(A)** Binding modes of the antagonist naloxone (cyan) compared to the agonist morphine (magenta) illustrated as in Fig. 2. **(B)** Analyzed binding modes were plotted based on the fingerprint analysis as shown in Fig. 2. The fingerprint of naloxone joins the group of the ligands activating upon depolarization (blue). **(C)** Average (mean ± SEM) FRET-based single cell recording of MOR-induced Gα_i_ activation under voltage clamp conditions is plotted for naloxone with the applied voltage protocol indicated below (n=7). **(D)** Voltage dependence of naloxone (blue) induced Gα_i_ activation was compared to morphine (magenta). The activation was determined by clamping the membrane from -90 mV to different potentials and plotted relatively to 0 mV. The data was fitted to Boltzmann function resulting in z-factor of 1.17 for naloxone and 0.8 for morphine and a V_50_-value of 31 mV for naloxone and -29 mV for morphine. **(E)** Representative recording of inward K^+^ currents in HEK293T cells expressing MOR and GIRK channels were the GIRK currents were evoked by naloxone and DAMGO. The currents were measured at -90 mV (depicted as blue dotted line) or at -20 mV (depicted as magenta dotted line). GIRK channels were blocked with 500 µM Ba^2+^ as indicated. Determination of activation level induced by naloxone is indicated by the filled blue box (or magenta box, respectively) compared to the activation induced by DAMGO (empty box) (as described before (Ruland et al., 2020)). **(F)** The GIRK current response evoked by naloxone was normalized to maximum response evoked by DAMGO at respective membrane potential. The responses at -90 and -20 mV were compared in the same recording, indicating an increased naloxone-induced current at -20 mV (p < 0.05, paired, two-tailed t-test).

All in all, this confirms the strong agonist specific voltage effect on the MOR, which is even able to convert antagonists to agonists. All the effects seem to be determined by the interaction pattern of each ligand, as different predicted binding modes – either between different ligands or of one ligand in a mutant receptor - are correlated with the extent and direction of the voltage effect.

## Discussion

In this study, we analyzed the binding modes of several clinically relevant opioid ligands by molecular docking calculations and subsequent experimental validation of the predicted binding modes by FRET-based functional signaling assays. We identified different predicted interactions for morphinan ligands versus methadone and fentanyl. These differential interaction patterns were connected to ligand-specific voltage sensitivity of the MOR. Furthermore, we were able to identify important regions in the receptor which we correlated with the voltage effect on the MOR.

Specifically, our molecular docking studies and subsequent fingerprint analysis, which calculated the interactions between a ligand and a receptor, revealed that morphine (or agonists with the morphinan scaffold) interacted with D147^3.32^, Y148^3.33^ and the water networks between Y148^3.33^ and H297^6.52^ as described before (Huang et al., 2015; Kapoor et al., 2020; Lipiński et al., 2019; Vo et al., 2021). Moreover, morphine displayed several interactions with helix 6, which were mainly missing for methadone and fentanyl, also consistent with previous findings (Kapoor et al., 2020; Lipiński et al., 2019). The observed binding pose for methadone indicated a salt bridge with D147^3.32^ as only direct interaction, comparable to the findings of Kapoor et al. For fentanyl, we identified an H-bond with Y326^7.43^ as critically important interaction. Indeed, we observed a strong right-shift of 4 orders of magnitude in the concentration-response curve for G_i_ protein activation, indicating a severe loss in potency for the tested Y326^7.43^ mutant, perfectly in line with our docking calculations. After further sampling of the fentanyl pose, we observed a salt bridge with D147^3.32^ for the best scored poses. This interaction was also seen by Vo et al. in MD simulations. However, the Y326^7.43^-interaction was missing in their simulation. Moreover, they postulated a second binding mode with H297^6.52^ as main interaction. This discrepancy can most likely be attributed to the fact that Vo et al. left out the two water molecules - which were important for ligand placement in our docking calculations - during their docking attempts. Lipiński et al., who included the water molecules, obtained results in their docking analysis that were comparable to our findings, underlining the importance of the water molecules for the calculations. However, the additional interaction with D147^3.32^, which is postulated to be crucial for fentanyl (Surratt et al., 1994; Weltrowska et al., 2010), only occurred after increased sampling of the conformational space around the docked pose of fentanyl. This suggests that fentanyl might adopt different stable binding modes and interactions, as also seen by Vo et al. In summary, with our approach we were able to corroborate the calculated binding modes experimentally by mutations or by the evaluation of the voltage sensitivity of the evaluated ligand.

We further evaluated several opioids in regard to their voltage sensitivity by means of FRET under conditions of whole cell voltage clamp. We identified ligands showing a strong increase in receptor activation upon depolarization of the membrane potential in a physiological range (morphine, buprenorphine, pethidine, etorphine, DAMGO, tramadol, PZM21 and naloxone). In contrast, other ligands displayed a decrease in activation (methadone, fentanyl, TRV130, loperamide and meptazinol). Met-enkephalin (Ruland et al., 2020) and SR17018 displayed no apparent voltage sensitivity. This opposite direction of the voltage effect cannot be explained by the differentiation between partial and full agonists. Both partial and full agonists were included in each of the tested groups. Moreover, biased agonists (PZM21, TRV130 and SR17018) were present in all groups. In conclusion, this indicated that the increased or decreased activation due to depolarization is not dependent on the degree of receptor activation. Additionally, the voltage effect was able to turn the antagonist naloxone into an agonist, comparable to the effects investigated for the P2Y_1_ receptor (Gurung et al., 2008).

Importantly, we detected the grouping of the opioids according to the direction of their voltage effect matched to a very high degree with the grouping based on the analysis of the fingerprints describing the docked binding poses. These results revealed a strong ligand specific voltage sensitivity, which seemed to be determined by the specific binding mode, and thus interaction pattern, of the ligands. Further analysis of the distinct interaction motifs of the ligand groups indicated three main interaction motifs determining the voltage effect. Helix 6 (W293^6.48^, H297^6.52^ and V300^6.55^) was indicated as important interaction site for the ligands which had an activating effect upon depolarization. In contrast, a motif located mainly on helix 3 (C217^45.50^, I144^3.29^, Y148^3.33^ and M151^3.36^) appears to be important for the ligands displaying a decrease in activation. Finally, a motif on helices 2, 3 and 7 (Q124^2.60^, N127^2.63^, D147^3.32^ and Y326^7.43^) was important for all ligands. A strong influence on ligand-specific voltage sensitivity of helices 3 and 6 was also reported for the muscarinic M_3_ receptor (Rinne et al., 2015). In general, there is still a lot of speculation about a possible general voltage sensing mechanism for GPCRs (Barchad-Avitzur et al., 2016; Hoppe et al., 2018; López-Serrano et al., 2020; Vickery et al., 2016). In this context, the involvement of a sodium ion bound to a conserved D was discussed (Vickery et al., 2016). This sodium ion seems to be important for the activation of the MOR (Selley et al., 2000; Sutcliffe et al., 2017). However, it was recently shown that this sodium ion is not involved in the voltage sensing mechanism of GPCRs (Hoppe et al., 2018). Our approach of combining in silico and in vitro methods enabled us to influence the binding mode of the opioids by site-directed mutagenesis and test the influence of the binding mode on voltage sensitivity. Overall, we were not able to change the directionality of the voltage effect on MOR activation for morphinan compounds. In contrast, for methadone and fentanyl we were able to change the direction of the voltage effect. Exchange of amino acids located in helices 3 and 6 displayed the largest effects on voltage sensitivity. Especially mutation of Y148^3.33^ resulted in an increased receptor activation upon depolarization for all tested ligands. A similar effect was induced by the H297^6.52^A mutation. For methadone, this led to a changed binding mode, introducing an interaction with Y148^3.33^, and resulting in an increased activation upon depolarization. It can be speculated that if the ligands are located closer to helix 3, the movement of helix 6, which is known to move outward upon receptor activation (Huang et al., 2015), could be increased upon depolarization. On the one hand, there could simply be more space for this movement if the ligands strongly interact with helix 3, further increasing the activation of the receptor. Supporting this hypothesis, we previously showed that the voltage effect induced by activation with morphine is primarily due to an increase of efficacy in receptor activation and not in affinity for the receptor (Ruland et al., 2020). On the other hand, ligands not showing this strong interaction with helix 3, such as fentanyl, could lose affinity for the receptor due to this movement or they might impede this movement, stabilizing the receptor in a more inactive state. What contradicts this hypothesis is that the Y326^7.43^F mutation, which, based on the docking calculations, altered the binding mode of fentanyl so that it interacted with Y148^3.33^, did not affect the voltage effect induced by fentanyl, but did change the direction of the voltage effect for methadone. It is possible that fentanyl can adopt different binding modes, as described by Vo et al. as well, as we also saw a possible interaction of the charged amine with D147^3.32^.

These results suggest that ligand-specific voltage sensitivity of MOR activation is mechanistically based on the interaction patterns between ligands and the receptor. Therefore, we propose that depolarization influences the conformation (or probability to reach certain conformations) of MOR in a way that increases the probability to activate receptors for ligands primarily interacting with helix 6 and decreases it for those ligands interacting with motif 2 in helix 3. In light of the observed ligand specific voltage sensitivity also seen with other receptors, this hypothesis might well apply to those receptors as well, if not to those for which voltage sensitivity has not been described yet. Our approach, strongly involving the opportunities enabled by in silico methods, allows the screening of a large number of predicted interactions and helps to choose the most information-rich receptor mutants and ligands for the subsequent in vitro analysis in a systematic and rational way. The MOR, with its diverse voltage pharmacology, was a good model system to illustrate the potential of this approach.

As MOR is expressed in neuronal tissue, which is highly excitable, a pharmacological relevance of voltage sensitivity of the MOR is very likely, albeit difficult to proof. We have already shown that the voltage sensitivity of the MOR is also reflected in brain tissue (Ruland et al., 2020). The voltage effect of the distinct opioid ligands might be taken into consideration as determinant of the clinical profile of opioids. A better understanding of the voltage dependence of the MOR, as achieved in our study, can potentially help with the development of safer and more effective opioids. It is known that neurons sensing pain depolarize more often. Development of opioid ligands with a voltage dependence stronger than morphine could therefore potentially act predominantly in these depolarized cells. This would be a novel way of precise drug targeting, possibly reducing side effects, which are still the main problems of opioid therapy.

## Acknowledgements

We thank Stefan Schulz and Andrea Kliewer (University of Jena, Germany) for the provision of the biased compounds PZM21, SR17018 and TRV130.

## Author contributions

S. B. K.: designed, performed and analyzed cellular experiments, designed, performed and analyzed the molecular docking calculations, performed and analyzed the fingerprint analysis and PCA, wrote the manuscript; V. J. Y. L.: designed the fingerprint analysis and PCA, performed and analyzed fentanyl sampling, reviewed and edited the manuscript; J. G. R.: designed, performed and analyzed cellular experiments, reviewed and edited the manuscript; P. K.: designed and supervised the molecular docking calculations and analyzed the data, reviewed and edited the manuscript; M. B.: wrote the manuscript, designed experiments and supervised the study.

## Competing interests

The authors declare no competing interests.

Supplemental Information is available for this paper.

## Materials and Methods

### Molecular docking and fingerprint analysis

The crystal structure of the active-state MOR (PDB code 5C1M (Huang et al., 2015)) was prepared for docking by deletion of the N-terminus up to residue 63 and the inclusion of two water molecules (HOH 505 and HOH 526). These were selected after a comparison of the distribution of all water molecules in the available crystal structures of opioid receptors (PDB codes: 5C1M, 4DKL, 4DJH, 4N6H) and analysis of their electron density including their EDIA-score (ProteinsPlus, ZBH, https://proteins.plus (Fährrolfes et al., 2017; Schöning-Stierand et al., 2020)). Using MakeReceptor (OpenEye Scientific Software, Santa Fe, NM, USA, http://www.eyesopen.com), the water molecules were defined as part of the receptor and D147^3.32^ (numbers in superscript are according to the Ballesteros-Weinstein enumeration scheme for GPCRs (Ballesteros and Weinstein, 1995)) as main interaction partner, as shown in (Surratt et al., 1994). Ligand preparation was performed with OMEGA (OpenEye, (Hawkins et al., 2010)), using isomeric SMILES from PubChem or ChEMBL. After docking buprenorphine, fentanyl, methadone and morphine using FRED (OpenEye, (Mc Gann, 2011)), the binding pocket and ligand of each protein-ligand complex was minimized with SZYBKI (OpenEye). The agonists were then compared in the different minimized binding pockets in order to select the optimally shaped receptor conformation for further calculations. The binding pocket of morphine was selected, as all agonists fitted without any steric clashes with the protein while satisfying most of their polar interaction possibilities. For each ligand, the top ranked 10-20 calculated poses were visually inspected (PyMol, Schrödinger, New York City, New York, USA, https://pymol.org) and the one or two poses that interacted most favorably with the MOR in terms of satisfied polar interactions in the absence of strained ligand geometry were selected for the fingerprint analysis. Sampling of fentanyl poses were performed with PyMol with “set_dihedral” function at five-degree intervals. Poses were minimized with OEtoolkit OEFF module (OpenEye) using steepest descent algorithm with MMFFAmber forcefield. The poses were ranked based on their minimized forcefield energy. The top 10,000 poses were clustered with CPPTRAJ (Roe and Cheatham, 2013) on the RMSD between heavy atom of the poses, stopping when five clusters are generated. Fingerprints were calculated using the program Arpeggio (Jubb et al., 2017), results were analyzed with principal component analysis using scikit-learn (Pedregosa et al., 2011) and the first two principal components were plotted. In that way, the 10 most important interactions were determined for each component. This procedure was repeated with the ligands split into two groups (defined based on their voltage sensitive behavior or their predicted voltage sensitive behavior). For comparison and analysis of the mutations, point mutations were inserted in the receptor by manually replacing and locally minimizing the selected amino acid with MOE (Chemical Computing Group, Molecular Operating Environment, Montreal, Canada). Then the docking and the following analysis were repeated as described before.

### Plasmids

cDNAs for rat MOR-wt, MOR-sYFP2, Gα_i_-YFP, Gα_o_-YFP, Gβ_1_-mTur2, Gγ_2_-wt, arrestin3-mTur2, GRK2-wt, GRK2-mTur2, Gα_i_-wt, Gβ_1_-wt, Gγ_2_-wt, bicistonic plasmid expressing GIRK3.1 and GIRK3.4 subunits and pcDNA3-eCFP have been described previously (Ruland et al., 2020). Gß-2A-cpV-Gy_2_-IRES-Gα_i2_- mTur2 was purchased from Addgene (Watertown, Massachusetts, USA, plasmid #69624 (van Unen et al., 2016)). Point mutations were introduced into MOR by site-directed mutagenesis and were verified by sequencing (Eurofins Genomics, Ebersberg, Germany). The following mutagenesis primers were used (sequence 5’→3’): Q124^2.60^E agtacactgccctttgagagtgtcaactacctg; N127^2.63^L cctttcagagtgtcttatacctgatgggaacatg; I144^3.29^S ctctgcaagatcgtgagctcaatagattactac; I144^3.29^V ctctgcaagatcgtggtctcaatagattactac; Y148^3.33^A gtgatctcaatagatgcctacaacatgttcacc; Y148^3.33^F cgtgatctcaatagatttctacaacatgttcaccag; M151^3.36^A atagattactacaacgcgttcaccagcatattc; K233^5.39^A ctgggagaacctgctcgcaatctgtgtctttatc; K233^5.39^E ctgggagaacctgctcgaaatctgtgtctttatc; V236^5.42^N cctgctcaaaatctgtaactttatcttcgctttc; W293^6.48^F gtatttatcgtctgctttacccccatccacatc; H297^6.52^A ctgctggacccccatcgccatctacgtcatcatc, H297^6.52^F ctgctggacccccatcaagatctacgtcatcatc; V300^6.55^A cccatccacatctacgccatcatcaaagcgctg; V300^6.55^F cccatccacatctacttcatcatcaaagcg, V300^6.55^L cccatccacatctacctcatcatcaaagcg; V300^6.55^N cccatccacatctacaacatcatcaaagcgctg; H319^7.36^Y cagaccgtttcctggtacttctgcattgctttgg; Y326^7.43^F gcattgctttgggtttcacgaacagctgcctg. The mutations Q124^2.60^E (Fowler et al., 2004), Y148^3.33^F (Xu et al., 1999), H297^6.52^A (Mansour et al., 1997; Spivak et al., 1997), H297^6.52^F (Spivak et al., 1997), H319^7.36^Y (Ulens et al., 2001) and Y326^7.43^F (Mansour et al., 1997) were evaluated before. As we performed our FRET and electrophysiological experiments in individual single cells, we did not test the expression levels of the differently mutated receptor versions, as the levels vary between every cell. To therefore exclude falsified conclusions on effects of the receptor mutants, we recorded concentrations response curves for every receptor variant and every used agonist.

### Cell Culture

All experiments in this study were carried out in HEK293T cells. Cells were cultured in high-dose DMEM supplemented with 10 % FCS, 2 mM L-glutamine, 100 U/ml penicillin and 0.1 mg/ml streptomycin at 37°C and 5% CO_2_. Cells were transiently transfected in 6 cm Ø dishes using Effectene Transfection Reagent according to manufacturer’s instructions (Qiagen) two days before the measurement. For MOR-induced Gα_i_ activation measurement, cells were transfected with 1 µg of MOR-wt or mutated MOR and 1 µg Gß-2A-cpV-Gy2-IRES-Gai2-mTur2, for measurements of voltage dependence of morphine fitted to Boltzmann function (Fig. 6), cells were transfected with 0.5 µg MOR-wt, 1 µg Gα_i_-YFP, 0.5 µg Gβ_1_-mTur2 and 0.25 µg Gγ_2_-wt. For measurement of MOR-induced GIRK currents, cells were transfected with 0,3 µg MOR-wt, 0.5 µg GIRK3.1/3.4 and 0.2 µg pcDNA3-eCFP. For MOR-induced Gα_o_ activation measurement, cells were transfected with 0.5 µg of MOR-wt, 1 µg G_o_-YFP, 0.5 Gß_1_-mTur and 0.25 µg Gy_2_-wt. For MOR-induced arrestin interaction, cells were transfected with each 0.7 µg of MOR- sYFP2, arrestin3-mTur2 and GRK2-wt. Cells were split on poly-L-lysine coated coverslips the day before the measurement. For MOR-induced GRK interaction, cells were transfected with 0.6 µg MOR-sYFP2, 0.6 µg GRK2-mTur2, 0.7 µg Gα_i_-wt, 0.6 µg Gβ_1_-wt and 0.6 µg Gγ_2_-wt

### Reagents

DMEM, FCS, penicillin/streptomycin, L-glutamine and trypsin-EDTA were purchased from Capricorn Scientific (Ebsdorfergrund, Germany). DAMGO acetate salt, buprenorphine-HCl, fentanyl citrate, tramadol-HCl and BaCl_2_ were purchased from Sigma-Aldrich (St. Louis, Missouri, USA). Etorphine-HCl (Captivon98©) was obtained from Wildlife Pharmaceuticals through Chilla CTS GmbH (Georgsmarienhütte, Germany). Loperamide-HCl was purchased from J&K chemicals (San Jose, CA, USA), meptazinol-HCl was purchased from Biozol (Eching, Germany), morphine hydrochloride was purchased from Merck (Darmstadt, Germany) and naloxone-HCl was purchased from Cayman Chemical (Ann Arbor, Michigan, USA). L-methadone-HCl and pethidine-HCl were purchased from Hoechst AG (Frankfurt, Germany). PZM21, SR17018 and TRV130 were a kind gift from Stefan Schulz and Andrea Kliewer, University of Jena, Germany (Gillis et al., 2020; Miess et al., 2018).

### FRET and electrophysiological measurements

Single-cell FRET measurements with or without direct control of the membrane potential were performed as described previously (Ruland et al., 2020). Using an inverted microscope (Axiovert 135, Zeiss) and an oil-immersion objective (A-plan 100x/1.25, Zeiss), CFP was excited by short light flashes of 430 nm (Polychrome V light source), fluorescence emission of YFP (F_535_) and CFP (F_480_) were detected by photodiodes (TILL Photonics Dual Emission System) with a sample frequency of 1 Hz, recording of data was performed with PatchMaster 2×65 (HEKA) and the FRET emission ratio of F_YFP_/F_CFP_ was calculated. After a necessary technical update of the setup, excitation was performed at 436 nm with a LED light source (precisExcite-100, 440 nm, CoolLED), and emission of YFP and CFP were split by an optosplit (Chroma) and detected with a CCD camera (RETIGA-R1, Teledyne Photometrics) and stored with VisiView software (Visitron Systems). As all measurements were normalized to a maximal answer within every measurement, the data was comparable between the two setup configurations. During measurements, cells were continuously superfused with either external buffer (137 mM NaCl, 5.4 mM KCl, 2 mM CaCl_2_, 1 mM MgCl_2_, 10 mM HEPES, pH7.3) or external buffer containing agonist in the respective concentration using a pressurized fast-switching valve-controlled perfusion system (ALA Scientific) allowing a rapid change of solutions. For FRET measurements under direct control of the membrane potential, cells were simultaneously patched in whole-cell configuration with the membrane potential set to a defined value by an EPC-10 amplifier (HEKA). For this, borosilicitate glass capillaries with a resistance of 3-7 MΩ were filled with internal buffer solution (105 mM K^+^-aspartate, 40 mM KCl, 5 mM NaCl, 7 mM MgCl_2_, 20 mM HEPES, 10 mM EGTA, 0.025 mM GTP, 5 mM Na^+^-ATP, pH 7.3). For measurement of GIRK currents, cells were measured in whole cell configuration analogue to the FRET measurements in 1 kHz sampling intervals with holding potentials of -90 or -20 mV, as indicated. As inward currents were measured, the used extracellular buffer was a high K^+^ concentration containing buffer (as external buffer, but with 140 mM KCl and 2.4 mM NaCl). All measurements were performed at room temperature.

### Data analysis and Statistics

FRET measurements were corrected for photobleaching (using OriginPro 2016) and were normalized to maximum responses within the same cell and recording. Further data analyzation was performed with GraphPad Prism 7 (GraphPad Software). Data is always shown (if not indicated differently) as mean ± SEM and group size defined as n. Statistical analysis were performed with a paired Student’s t-test or a two-tailed unpaired t-test with Welch’s correction (as normality of data distribution wasn’t given for every group) or, for more than two groups, by an ordinary one-way ANOVA (as SD’s were significantly different a Brown-Forsythe and Welch’s ANOVA test were performed) with Dunnet’s T3 multiple comparisons test, as indicated. Differences were considered as statistically significant if p ≤ 0.05. Concentration-response curves were fitted with a non-linear least-square fit with variable slope and a constrained top and bottom using following function:

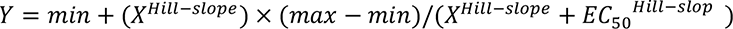

where min and max are the minimal and maximal response and EC_50_ the half-maximal effective concentration. Voltage sensitive behavior was analyzed by normalizing the answer at +30 mV (mean of last 10 s before repolarization) to the answer at -90 mV (mean of last 10 s before depolarization) with previous normalization of the whole trace to the agonist-induced answer at -90 mV as max. response. For analyzation of charge movement and V_50_-values, answers were subtracted for -90 mV and normalized to 0 mV. These values, now normalized for the degree of receptor activation (R) reflected by Gα_i_ activation, were fit to a single Boltzmann function. The equation used for fitting was

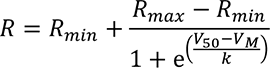

where R_min_ and R_max_ were the minimal and maximal response, V_M_ the respective membrane potential, V_50_ the voltage of half-maximal effect on Gα_i_ activation and k the slope factor. For calculation of the z factor, the net charge movement upon change in V_M_ across the membrane, following equation was used:

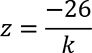

For analyzation of GIRK current response evoked by naloxone, the responses to naloxone at either -90 or -20 mV were normalized to the max. response evoked by DAMGO at the respective V_M_ and values generated in the same recording were compared.

## Supplemental Information

**Figure S1:**
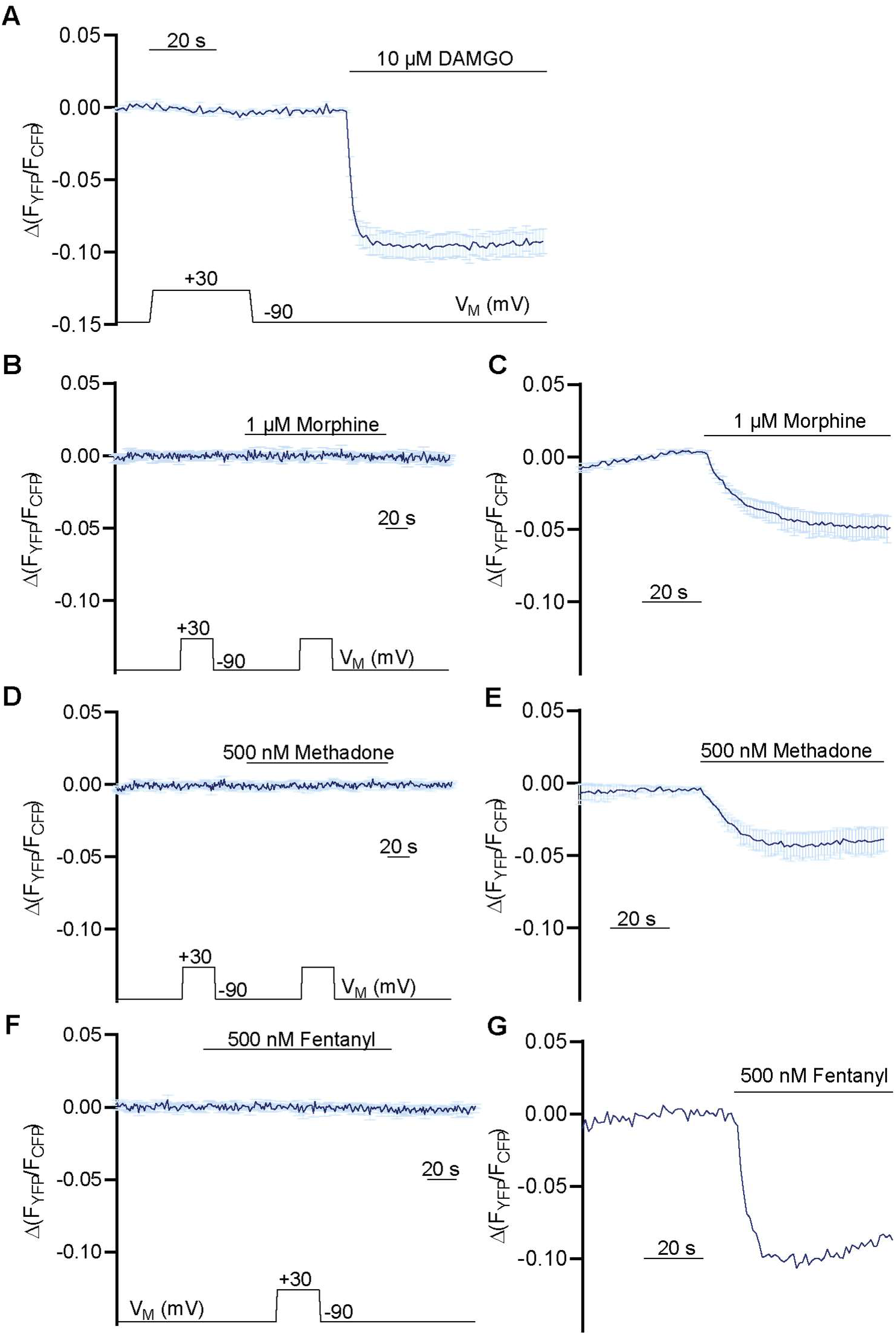
Control measurements for voltage effect of MOR in Gα_i_ activation assay. **(A)** Averaged FRET-based single cell recordings of MOR-induced Gα_i_ activation under voltage clamp conditions with WT receptor in HEK293T cells (mean ± SEM; n=8). The applied voltage protocol is indicated below. Depolarization during application of buffer and without application of agonists has no effect. **(B, D, F)** Averaged FRET-based single cell recordings of Gα_i_ activation under voltage clamp conditions without transfection of the MOR receptor in HEK293T cells (mean ± SEM B: n=7, D: n=7, F: n=6). Neither a depolarization under application of agonist nor the depolarization under applications of agonist showed an effect. **(C, E)** Averaged FRET-based single cell recordings of Gα_i_ activation induced by MOR wt in HEK293T cells (mean ± SEM, E: n=4, E: n=4). Measurements were performed in parallel to the experiments depicted in B and D as positive control. **(G)** Representative FRET-based single cell recording of Gα_i_ activation induced by MOR wt in HEK293T cells. Measurements were performed in parallel to the experiments depicted in F as positive control.

**Figure S2:**
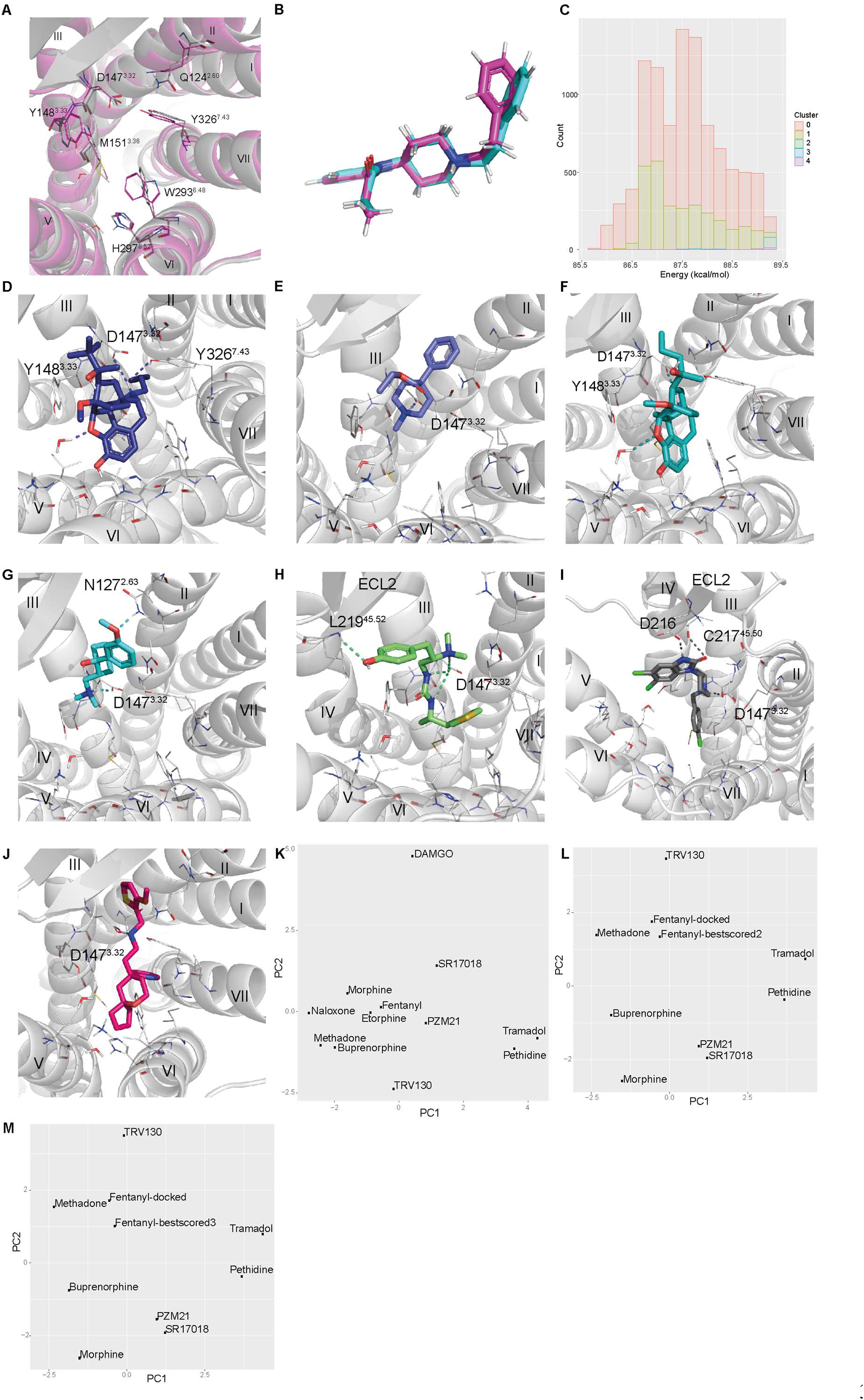
Binding modes of different opioids docked to MOR. **(A)** Comparison of the binding pocket used for our docking calculations (grey, based on the crystal structure of the active MOR, PDB: 5C1M) and the cryo-EM structure of DAMGO bound to MOR (PDB: 6DDE). Both conformations differ only slightly in the position of W293 and Q124. **(B)** Orientation of the representatives of the two most populated clusters of fentanyl which were discernable out of the top 10.000 poses. Cluster representative one is depicted in magenta, cluster representative two is depicted in cyan. **(C)** Clusters of the top 10.000 poses plotted against the energy values after minimization. **(D-J)** The binding modes of different opioids were analyzed regarding their fingerprints in Fig. 2D. The fingerprints were calculated based on the binding poses of buprenorphine (D), pethidine (E), tramadol (F), PZM21 (G), SR17018 (H), loperamide (I) and TRV130 (J). **(K)** The binding mode of DAMGO presented in 6DDE was aligned to the conformation used for our docking calculations and the fingerprints were analyzed. As DAMGO is a large peptide, the fingerprint differs substantially from the other evaluated opioid ligands. **(L-M)** Different conformers of fentanyl were sampled at three different dihedral angels and ranked according to the minimized energy. The calculated fingerprint of the second (L) and third (M) best scored conformer showed just a slight difference between the docked fentanyl (Fig. 2C and F) and the best scored fentanyl conformer (Fig. 2D and F).

**Figure S3:**
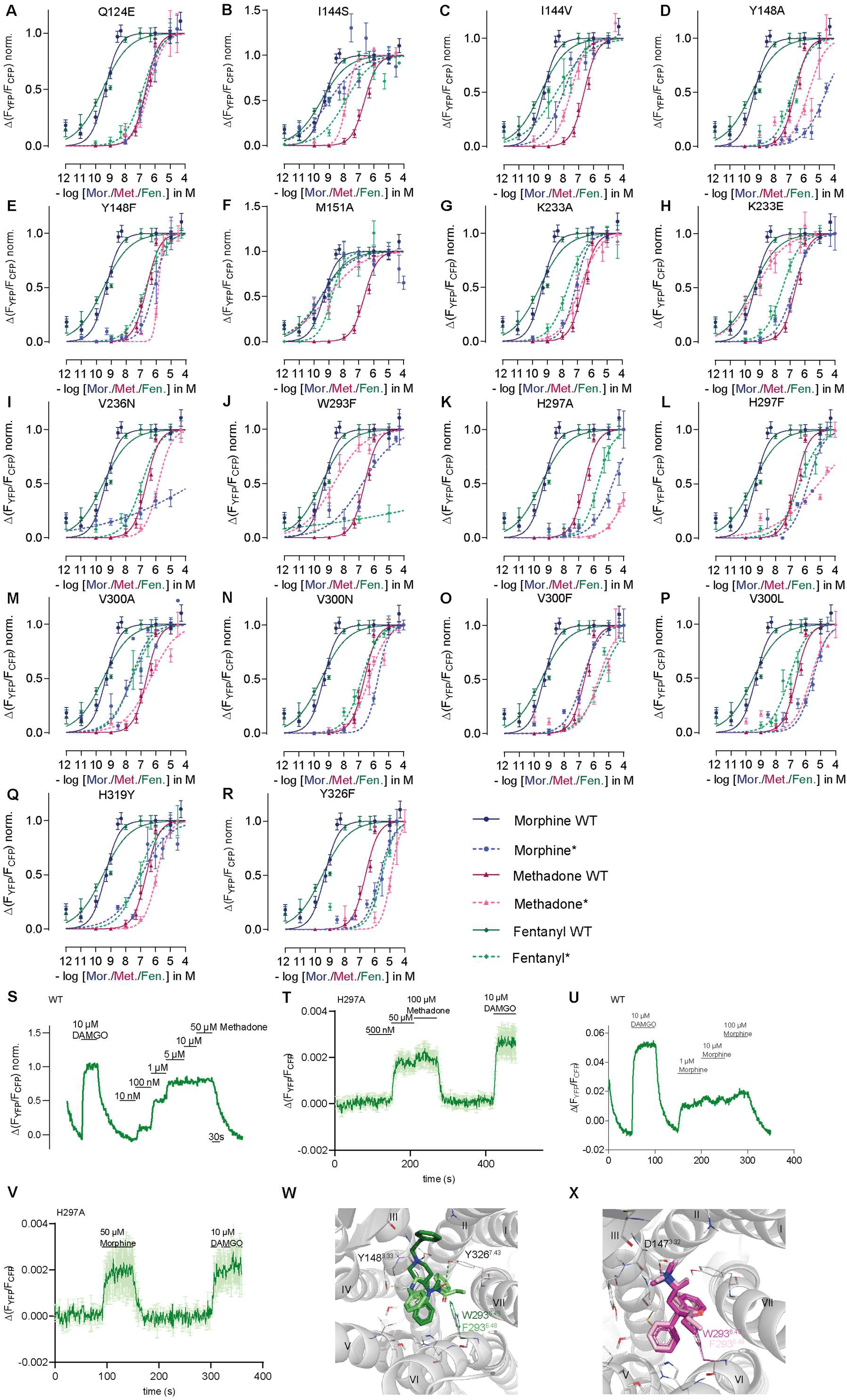
Functional effects of the insertion of mutations displayed by Gα_i_ activation and GRK2 interaction. **(A-R)** Concentration-response curves for Gα_i_ activation induced by mutated versions of MOR measured by single-cell FRET. Cells expressing MOR WT or mutated versions of the receptor were stimulated with morphine (blue), methadone (magenta) or fentanyl (green). Mutated versions of receptor are shown as dotted line. Data shown as mean ± SEM. For simplification, maximum Gα_i_ activation induced by the respective agonist is set as 1. EC_50-_values were calculated, normalized to WT and plotted in Fig. 3I. **(S, U)** Representative FRET-based single cell recording of MOR-GRK2 interaction induced by agonist application. Maximum activation for normalization was induced by 10 µM DAMGO. **(T, V)** Average single-cell recording of MOR-H297A mutant. For methadone (T), a saturation of the assay can be achieved with extremely high concentrations of methadone. For morphine (V), there’s an interaction is detectable as well. As the Gα_i_ activation displays a strong amplification, conclusions on efficacy changes induced by mutants can only be evaluated by direct one-to-one interactions like the GRK interaction. However, as the efficacy of activation induced by DAMGO seems to be weakened by this mutation, reliable efficacy values for the mutations can’t be calculated as there was no normalization possible. **(W, X)** Insertion of mutation W293^6.48^F changes binding mode of fentanyl (W) but doesn’t influence methadone (X).

**Figure S4:**
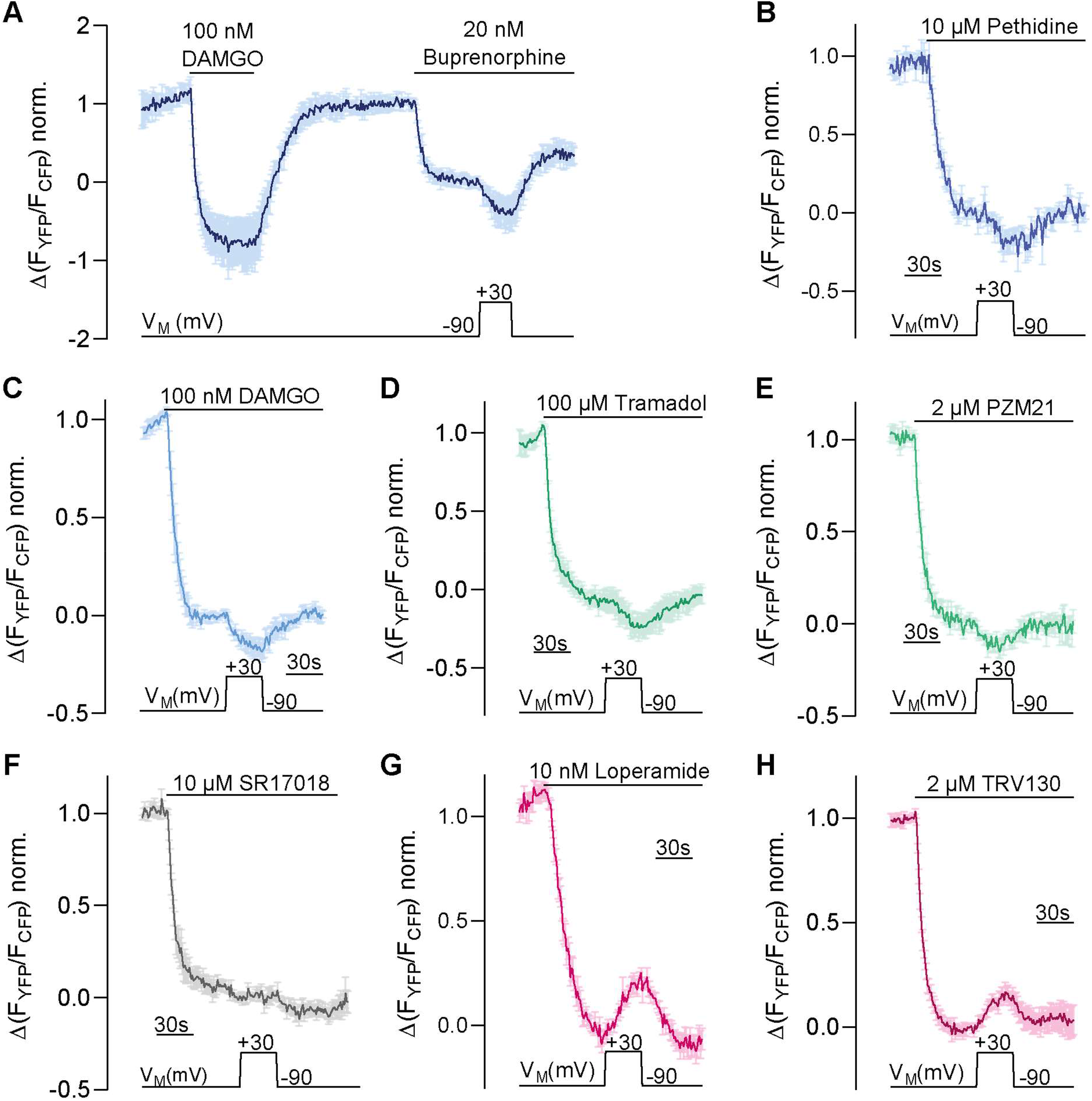
Agonist specific voltage sensitive behavior of the MOR. **(A-H)** Average FRET-based single cell recording of MOR-induced Gα_i_ activation under voltage clamp conditions plotted for the indicated agonists (mean ± SEM; A: n=6, B: n=6; C: n=13; D: n=5; E: n=6; F: n=6; G: n=9, H: n=7). The applied voltage protocol is indicated below. All agonist were applied at a non-saturating concentration inducing approx. same G_i_ activation level, as representatively indicated by the application of DAMGO in A.

**Figure S5:**
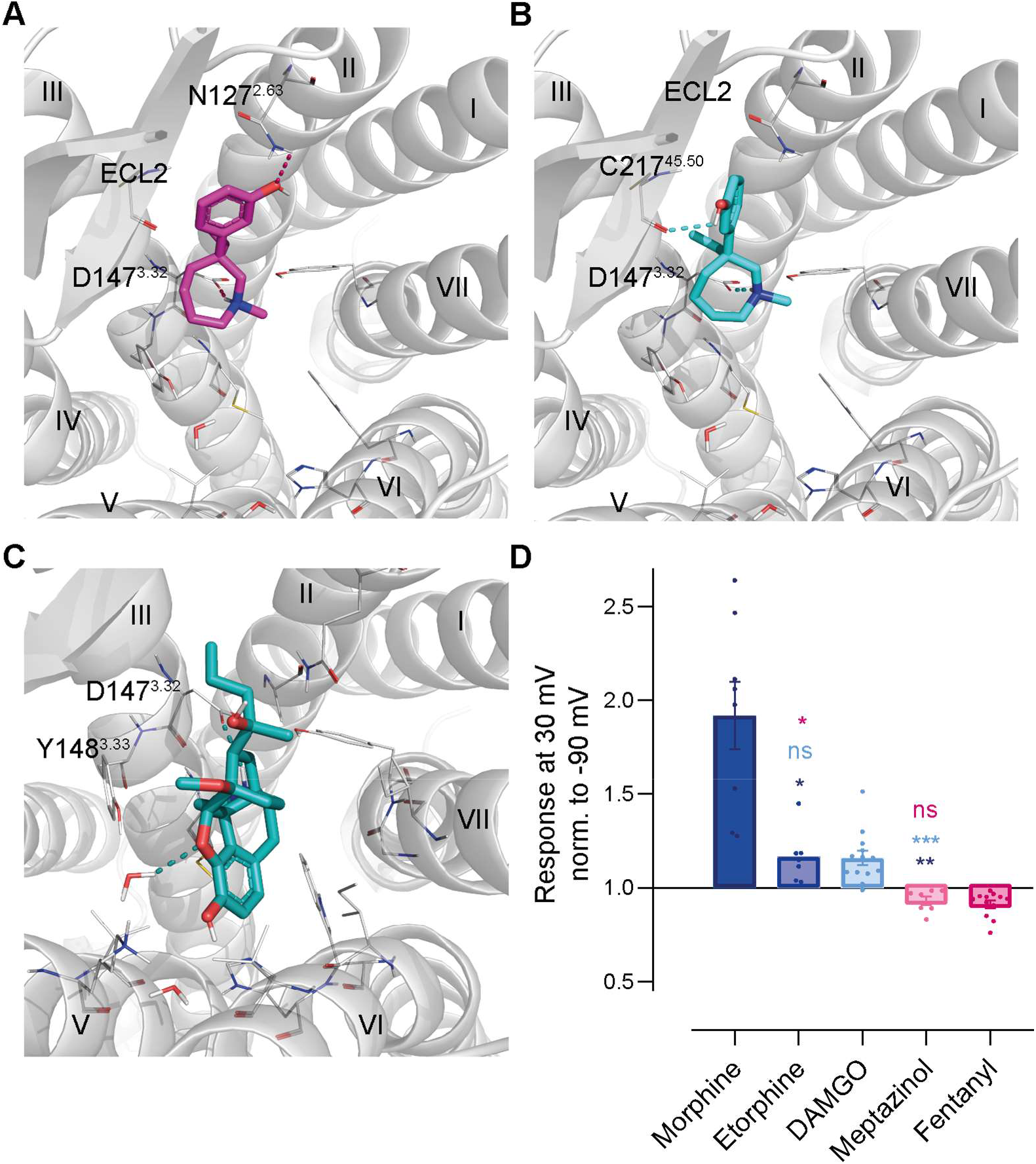
Binding modes and voltage effect of meptazinol and etorphine. **(A, B)** Meptazinol docked to MOR WT, in possible binding mode 1 (A) and binding mode 2 (B). **(C)** Etorphine docked to MOR WT. **(D)** Analysis of voltage sensitivity of MOR for meptazinol and etorphine of measurements shown in Fig. 5 A and E. The response of agonist-induced Gα_i_ activation at +30 mV was normalized to the response at -90 mV. Statistical significance was calculated compared to depolarization effect induced by morphine (dark blue), DAMGO (bright blue) and fentanyl (magenta) by an ordinary one-way ANOVA (p<0.0001) with Dunnett’s T3 multiple comparisons test (ns p>0.05, ** p<0.005, *** p<0.0005, **** p<0,0001).

**Figure S6:**
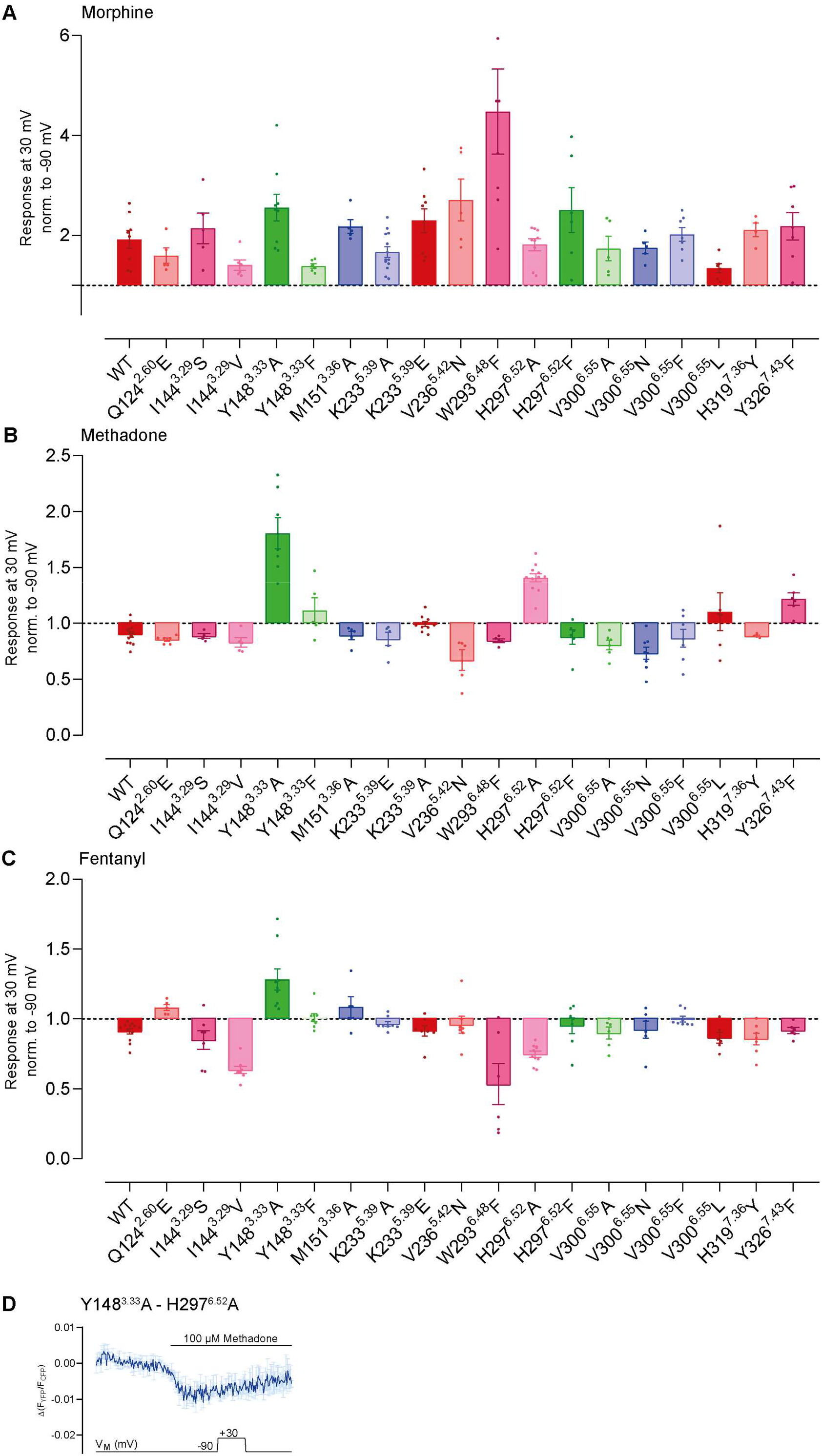
Altered binding modes influence voltage sensitivity of the MOR activated by morphine. **(A-C)** Average FRET-based single cell recording of MOR-induced Gα_i_ activation under voltage clamp conditions were measured as displayed in Fig. 6 A-D, analyzed and plotted in a bar graph regarding the inserted mutation and the induced voltage effect. Agonists (A: Morphine, B: Methadone, C: Fentanyl) were applied in a non-saturating concentration inducing approx. same G_i_ activation level, determined for every mutation in Fig. S3. **(D)** Average FRET-based single cell recording of MOR- induced Gα_i_ activation under voltage clamp conditions plotted for methadone with the double-mutant Y148A-H297A. The double mutation displays a very low activity, as there’s just a weak FRET-ratio change by extreme high methadone concentrations (mean ± SEM; n=4). For this reason, double mutations weren’t analyzed further.

**Figure S7:**
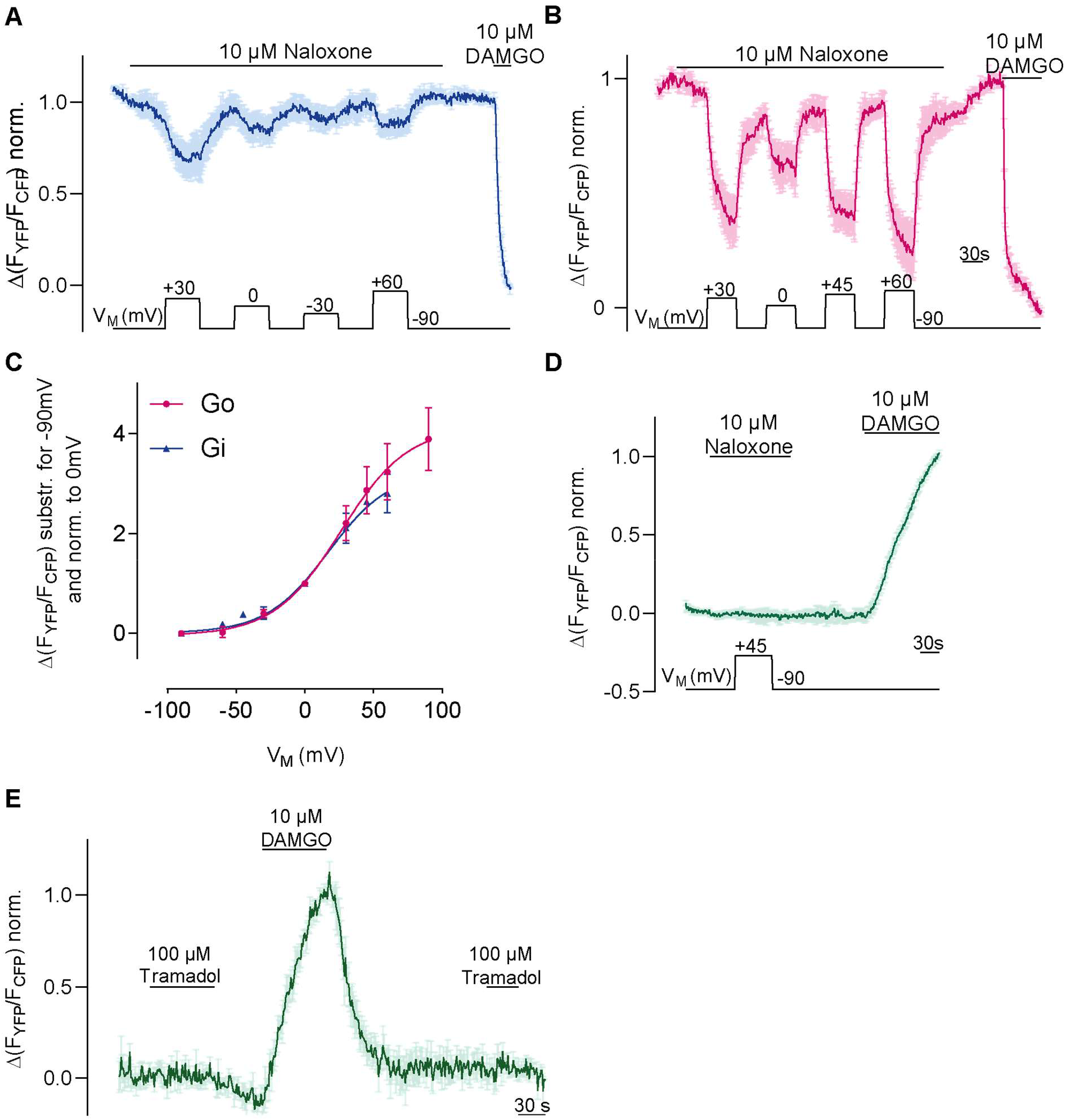
Depolarization converts the antagonist naloxone to an agonist. **(A)** Representative FRET-based single cell recording of MOR-induced Gα_i_ activation under voltage clamp conditions used for fit in Fig. 6d and S9c, the voltage protocol indicated below (mean ± SEM, n=4). **(B)** Representative FRET-based single cell recording of MOR-induced Gα_o_ activation under voltage clamp conditions used for fit in c, the voltage protocol indicated below (mean ± SEM, n=7). **(C)** Voltage dependence of naloxone-induced Gα_i_ activation (blue) compared to Gα_o_ activation (magenta). Activation was determined by clamping the membrane from -90 mV to different potentials and plotted relatively to 0 mV. Data was fitted to Boltzmann function resulting in z-factor of 1.17 for Gα_i_ and 1.2 for Gα_o_ and a V_50_- value of 31 mV for Gα_i_ and 27 mV for Gα_o_. **(D)** Average FRET-based single cell recording of arrestin-mTur2 interaction with MOR-sYFP2 under voltage clamp conditions, the voltage protocol indicated below (mean ± SEM, n=7). **(E)** Average FRET-based single cell recording of arrestin-mTur2 interaction with MOR-sYFP2 induced by the weak partial agonist tramadol (mean ± SEM, n=5).

